# Evidence of a conserved thermodynamic signature of growth state in living systems

**DOI:** 10.64898/2026.11.03.764118

**Authors:** Roberto Zivieri, John Reeves, Swetansu Pattnaik, Andre Martins Reis, Anthony Xu, David M. Thomas, Wilson H. Lee, Zhipeng Lu, Ira Deveson, Arcadi Cipponi

## Abstract

Living systems operate far from thermodynamic equilibrium, sustaining complexity through continuous free-energy influx and heat dissipation. Yet the higher-order functional constraints imposed by non-equilibrium dynamics remain largely unexplored.

To formalize how changes in cellular entropy production constrain intracellular organization across proliferative state transitions, we developed a stochastic-thermodynamic model linking reduced dissipative drive to transient increases in intracellular configurational entropy.

Within this framework, we examined abundance-weighted predicted RNA folding stability as a transcriptome-level projection of configurational remodelling.

Across the cell cycle, we observed coordinated, phase-dependent changes in the distribution of transcript-level RNA folding free energies.

Extending this analysis to stress adaptation, we find that human cancer cells exposed to molecularly targeted agents and nutrient-limited *Escherichia coli* converge on expressed RNA pools enriched for transcripts with less negative RNA folding free energies, despite marked heterogeneity in the transcriptional responses within each system. Opposing shifts during acquired resistance under continued drug exposure and bacterial recovery from starvation further support an association with growth state.

Together, these findings suggest that transcriptome organization extends beyond gene-regulatory programs to collective biophysical properties that may reflect non-equilibrium constraints shaping cellular-state transitions.

## Main

The stochastic components of evolution, together with the multiplicity of viable outcomes, can make the possible forms of molecular and cellular organization appear virtually limitless. Yet accumulating evidence indicates that the space of biological possibilities is tightly constrained. Recurrent design principles, functional convergence, and the restricted occupation of molecular morphospaces underscore how physical and chemical laws have consistently shaped the structure and dynamics of living systems^1–3^. Ultimately, these constraints are embodied in the physical basis of life, where cells operate as non-equilibrium open systems^4^ that sustain low internal entropy through continuous free energy influx and dissipation of metabolically generated heat. At the molecular scale, directionality arises from the coupling of thermodynamic forces to chemical and metabolic fluxes^5^.

To formalize thermodynamic constraints on cellular-state transitions, specifically cell-cycle progression and entry into growth arrest, we developed a coarse-grained stochastic model (see Supplementary Information for the full derivation). These transitions involve changes in metabolic and biosynthetic a ctiv ity that can alter both entropy production and export. In this model, the probability distribution *P*(Γ, *t*) over accessible intracellular configurations Γ, defines the configurational entropy:

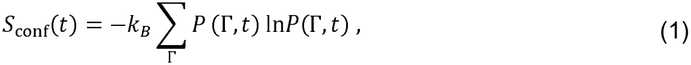

where *k*_*B*_ is Boltzmann’s constant and *t* denotes time. Configurational entropy evolves according to the balance equation:

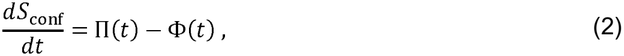

where the internal irreversible entropy-production rate Π(*t*), and net outward entropy-flow rate Φ(*t*) are defined by

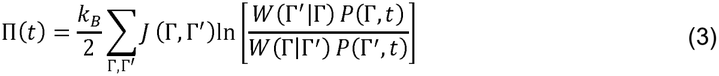

and

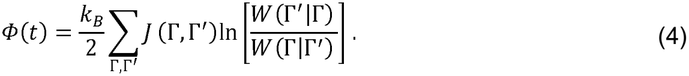

Here *W*(Γ’ | Γ) denotes the transition rate from configuration Γ to Γ’, and *J*(Γ, Γ’) is the corresponding net probability current. The outward entropy flow comprises contributions from heat dissipation and molecular exchange with external reservoirs

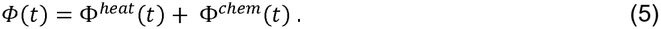

To model entry into growth arrest induced by cytostatic treatment or nutrient limitation, we treat the initial proliferative state as a nonequilibrium steady state, with Π^(0)^ ≃ Р^(0)^ > 0

Experimental metabolic evidence motivates the assumption that entropy export initially declines more strongly than internal entropy production^6–16^ (Supplementary Information, Sections 1.5.2-1.5.3). The entropy balance in Eq. (2) therefore implies a transient increase in intracellular configurational entropy:

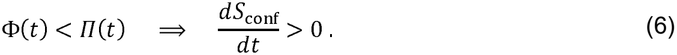

Beyond growth arrest, cell-cycle progression provides a complementary physiological setting, as successive phases differ in their metabolic and biosynthetic demands. These phase-dependent changes can alter the balance between entropy production and export, leading to increases or decreases in configurational entropy.

This framework provides the rationale for examining whether such configurational remodeling is reflected in molecular subsystems of the cell. Among these, the transcriptome is particularly informative, as gene expression responds rapidly to physiological and environmental cues^17^.

Each transcript carries sequence-encoded structural properties, characterized here by a length-normalized predicted folding free energy (Δ*G*_fold_), with more-negative values indicating greater predicted secondary-structure stability. This descriptor was calculated under fixed model conditions (Methods) and held constant across biological states.

Changes in relative transcript abundance can shift the folding-stability profile of the expressed RNA pool. The resulting shift in abundance-weighted mean folding free energy from reference state *r* to comparison state *c* is therefire given by

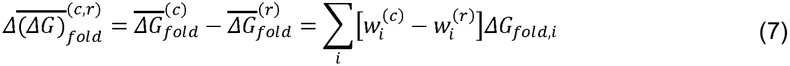

where 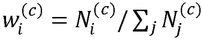 is the normalized abundance weight of transcript in state *c*, 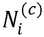 is its abundance, and *j* indexes all transcripts retained for the comparison.

We therefore asked whether systematic shifts in this collective biophysical property across proliferative-state transitions, could provide a transcriptome-level projection of the predicted configurational remodelling, with an expected shift towards a less negative abundance-weighted mean Δ*G*_fold_ during growth arrest.

To address this question, we first assessed the folding predictions against experimental data and then quantified RNA-pool reweighting across human and bacterial cell-state transitions in terms of the shift defined in Eq. (7). This revealed phase-dependent changes during cell-cycle progression and a convergent shift in the expected direction during growth arrest, despite divergent transcriptional responses.

These findings suggest a conserved biophysical dimension of transcriptome organization arising from abundance-driven reweighting of sequence-encoded RNA properties. This collective feature may reflect higher-order constraints imposed by non-equilibrium dynamics.

## Results

### RNA folding predictions are supported *in vivo* and reveal biotype- and pathway-specific stability profiles

To establish biological relevance, we first sought experimental support for the sequence-based folding predictions and then examined whether distinct RNA biotypes and transcripts associated with defined signaling and functional pathways display characteristic RNA secondary-structure stability profiles.

RNA molecules populate dynamic ensembles of alternative secondary structures^18^ that are, in principle, more fully described by partition-function-based approaches. However, transcriptome-wide analyses require a scalable and directly comparable descriptor. We therefore calculated the minimum-free-energy (MFE) secondary structures across 385,623 GENCODE v47 human transcripts using ViennaRNA^19^ under fixed folding conditions (Methods). For each transcript, Δ*G*_fold_ was defined as the predicted Gibbs free-energy change associated with folding the RNA into its MFE structure, normalized by transcript length and expressed in kcal·mol^−1^·nt^−1^ (Supplementary Table 1). More negative Δ*G*_fold_ values indicate greater predicted stability, whereas less negative values indicate reduced predicted stability.

To assess robustness, we compared ViennaRNA estimates with predictions generated by the RNAstructure^20^ algorithm (Supplementary Table 2). The two methods produced near-identical raw Δ*G*_fold_ estimates and transcript rankings (Pearson r = 0.9999; Fig.1A), and Bland–Altman analysis confirmed close agreement, revealing a minor shift towards more negative RNA structure estimates (mean difference = −1.84 kcal·mol^−1^; Fig. 1B).

**Fig. 1.**
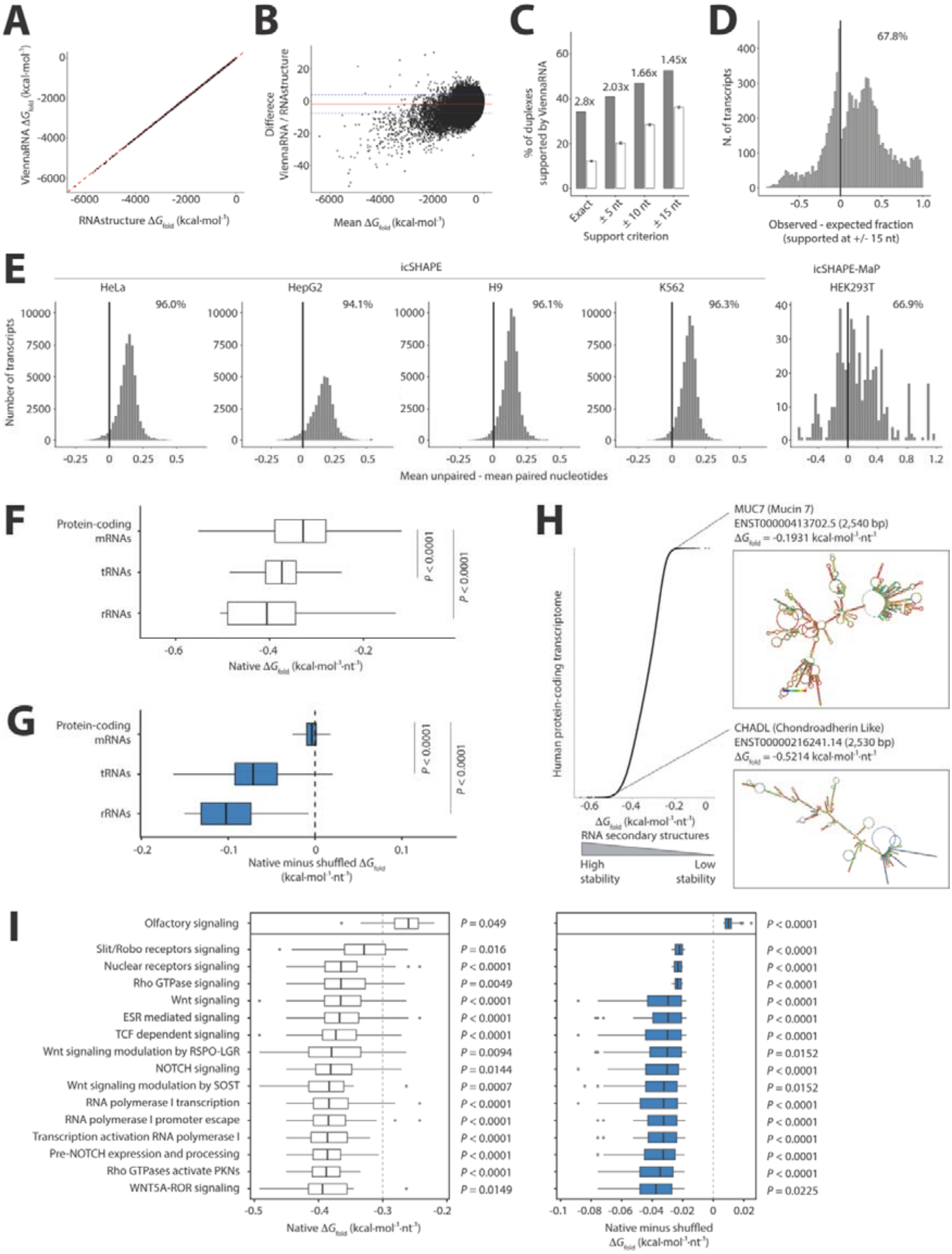
RNA folding predictions are supported *in vivo* and reveal biotype- and pathway-specific stability profiles. (**A**) Transcript-level Δ*G*_fold_ estimates from ViennaRNA and RNAstructure. Pearson r, Spearman ρ and linear-model R^2^ = 0.9999. (**B**) Bland–Altman comparison: solid red line, mean difference; dashed blue lines, 95% limits of agreement. (**C**) Fraction of 32,564 PARIS2 duplex groups from 9,330 transcripts supported by at least one ViennaRNA-predicted base pair using the unexpanded duplex-arm intervals (“Exact”) or relaxed ±5-, ±10-, and ±15-nucleotide criteria. Grey bars indicate observed support, and white bars indicate the mean support across 1,000 matched within-transcript randomizations; error bars show the 2.5th–97.5th percentiles of the random distributions. Values above the bar pairs indicate fold enrichment over the mean random expectation. One-sided randomization tests; *P* = 0.0009 for each criterion. (**D**) Distribution of transcript-level observed-minus-expected fractions of supported PARIS2 duplexes at ±15 nucleotides, using the matched within-transcript random model. The vertical black line marks zero; the percentage indicates transcripts with greater observed than expected support. (**E**) Nucleotide-resolution validation of ViennaRNA paired/unpaired assignments using *in vivo* icSHAPE and icSHAPE-MaP accessibility datasets. Histograms show the transcript-level mean accessibility difference between ViennaRNA-predicted unpaired and paired nucleotides. Positive values indicate higher experimental accessibility at predicted unpaired positions. Percentages denote the fraction of transcripts with positive mean unpaired-minus-paired accessibility differences in each dataset. (**F**) Distribution of normalized RNA Δ*G*_fold_ values across protein-coding mRNAs, tRNAs, and rRNAs. Wilcoxon rank-sum tests. (**G**) Differences between native and dinucleotide-preserving shuffled normalized Δ*G*_fold_ values across the same biotypes. Negative values indicate greater predicted stability of native sequences. Wilcoxon rank-sum tests. (**H**) Normalized Δ*G*_fold_ distribution across human protein-coding transcripts, with representative centroid structures for transcripts of similar length but contrasting predicted stability. (**I**) Native normalized Δ*G*_fold_ values (white bars) and native-minus-shuffled normalized Δ*G*_fold_ differences (blue bars) for genes belonging to ORA-enriched Reactome / KEGG pathways. *P* values are from ORA enrichment tests (left) and two-sided Wilcoxon signed-rank tests against zero (right), with BH correction.

Although primary RNA sequence constrains secondary-structure formation, the cellular RNA structurome is additionally shaped by RNA-binding proteins, helicase-mediated remodelling, RNA modifications and local ionic conditions^21^. We therefore assessed whether the sequence-encoded structural features predicted by ViennaRNA were supported by independent *in vivo* RNA structure-probing datasets.

We first analyzed PARIS2^22^ data from human HEK293T cells, in which RNA duplexes are represented as pairs of interacting transcript intervals rather than nucleotide-resolved base-pair maps. Accordingly, we compared intramolecular PARIS2 duplexes with ViennaRNA-predicted base pairs using the reported duplex-arm intervals (exact matching) and relaxed ±5-, ±10-, or ±15-nucleotide windows (Methods). Across 32,564 mapped PARIS2 duplexes from 9,330 transcripts, ViennaRNA predictions supported 34.1% of duplexes under exact base-pair matching, increasing to 52.6% with the relaxed ±15-nucleotide window. To account for chance overlap, we compared real duplexes with matched within-transcript random duplexes preserving transcript identity, arm length and inter-arm spacing. Real PARIS2 duplexes showed greater ViennaRNA support than the matched null expectation across all criteria, with enrichment ranging from 2.80-fold under exact matching to 1.45-fold at ±15 nucleotides (*P* = 0.0009 for each criterion; Fig. 1C). At the transcript level, support for real duplexes exceeded random expectation in 60.2% of evaluable transcripts under exact matching and 67.8% under the ±15-nucleotide criterion (Fig. 1D), indicating broadly distributed, statistically significant but partial concordance, consistent with the interval-based nature of this assay and its lack of complete nucleotide-resolved base-pair assignments.

We therefore extended these validations to nucleotide-resolution RNA structure-probing datasets, enabling direct comparison of ViennaRNA-predicted paired and unpaired states with *in vivo* nucleotide accessibility. We analyzed icSHAPE^23^ data from H9, HeLa, HepG2, and K562 cells, together with HEK293T icSHAPE-MaP^24^ data, in which higher reactivity scores indicate more accessible or conformationally flexible nucleotides, whereas lower scores indicate structurally constrained or base-paired positions. Following transcript-coordinate integration and coverage filtering (Methods), 41,875–62,934 transcripts were retained across the four icSHAPE datasets, together with 537 transcripts from the HEK293T icSHAPE-MaP dataset; joined-position and coverage summaries shown in Supplementary Fig. 1A–C.

Across all five datasets, nucleotides predicted to be unpaired displayed higher icSHAPE/icSHAPE-MaP reactivity than nucleotides predicted to be paired, yielding transcript-level accessibility differences that were significantly shifted toward positive values (*P* < 0.0001). Positive mean unpaired-minus-paired accessibility differences were observed in 96.1% of H9 transcripts (60,489/62,934), 96.0% of HeLa transcripts (52,677/54,887), 94.1% of HepG2 transcripts (39,407/41,875), 96.3% of K562 transcripts (56,877/59,047), and 66.9% of HEK293T transcripts (359/537; Fig. 1E). Coverage-stratified analyses confirmed that ViennaRNA experimental concordance was retained across both lower- and higher-coverage transcript subsets, indicating that the agreement was not driven by densely covered transcripts alone (Supplementary Fig. 1D). Concordance was also preserved after collapsing isoforms to Ensembl canonical transcripts, with positive mean accessibility differences observed in 82.6–96.8% of genes across datasets (Supplementary Fig. 1E).

Together, these data show that ViennaRNA-predicted paired and unpaired states are reflected in experimental icSHAPE/icSHAPE-MaP accessibility profiles across multiple human cell lines, providing broad *in vivo* support for ViennaRNA-derived folding metrics as scalable descriptors of sequence-dependent secondary-structure stability.

We next examined whether RNA biotypes exhibit structural-stability profiles consistent with their functions. RNA classes subject to stringent conformational constraints, such as tRNAs and rRNAs, are expected to exhibit highly stable secondary structures. More broadly, if sequence-encoded structural stability contributes to transcriptome organization, transcripts associated with distinct functional pathways should occupy non-random positions within the predicted Δ*G*_fold_ distribution.

Protein-coding transcripts showed significantly less negative normalized Δ*G*_fold_ values than tRNAs and rRNAs, which exhibited greater predicted stability (*P* < 0.0001 for both comparisons; Fig. 1F). To determine whether these differences reflected sequence-order-dependent stabilization rather than nucleotide composition alone, we compared native transcripts with matched dinucleotide-preserving shuffled controls (Methods). Native sequences were more stable than shuffled controls across all three RNA classes, but this excess stability was much greater for tRNAs and rRNAs than for protein-coding transcripts. Mean native-minus-shuffled values were −0.00577, −0.0668 and −0.0941 kcal·mol^−1^·nt^−1^ for protein-coding transcripts, tRNAs and rRNAs, respectively, with more than 93% of tRNAs and rRNAs showing greater native stability, compared with 69.0% of protein-coding transcripts (Fig. 1G). This pattern is consistent with structurally flexible regions facilitating mRNA translation, whereas compact, conserved tRNA and rRNA structures are required for codon recognition and ribosomal function.

To illustrate extremes of conformational stability among protein-coding transcripts, we selected transcripts of similar length from the less- and more-negative tails of the normalized Δ*G*_fold_ distribution and visualized their predicted centroid structures (Methods). The Mucin 7 (*MUC7*) isoform ENST00000413702.5, from the less-negative tail, showed sparse predicted base pairing, whereas the Chondroadherin-Like (*CHADL*) isoform ENST00000216241.14, from the more-negative tail, showed extensive predicted base pairing (Fig. 1H).

To move from single-transcript examples to a system-level analysis, we asked whether sequence-encoded RNA structural stability was non-randomly distributed across Reactome and KEGG pathways. Over-representation analysis (ORA) identified significantly enriched pathways with contrasting RNA structural-stability profiles. Transcripts contributing to olfactory signaling showed less-negative native Δ*G*_fold_ values, whereas those associated with developmental patterning, morphogenesis, cytoskeletal remodeling and proliferation, including WNT, Notch, RHO GTPase and ROBO/Slit signaling, showed more-negative values. Relative to matched shuffled controls, olfactory transcripts had lower predicted stability, whereas transcripts associated with growth and development had greater stability (Fig. 1I). These associations are consistent with the model’s prediction that increased proliferative activity is accompanied by greater predicted RNA secondary-structure stability (Supplementary Information).

Together, these findings provide *in vivo* support for sequence-based RNA structural predictions and show that stability differs among RNA biotypes and is associated with distinct functional pathways.

### Predicted RNA structural stability is coupled to transcriptome remodeling during cell-cycle progression

To extend these functional associations to dynamic cellular states, we focused on transcriptome remodeling across human cell-cycle phases (Fig. 2A). Metabolic and biosynthetic demands vary throughout the cell cycle: G1 supports protein synthesis and organelle biogenesis, S phase prioritizes nucleotide synthesis and DNA replication, and G2 is characterized by increased lipid biosynthesis for membrane generation^25^.

**Fig. 2.**
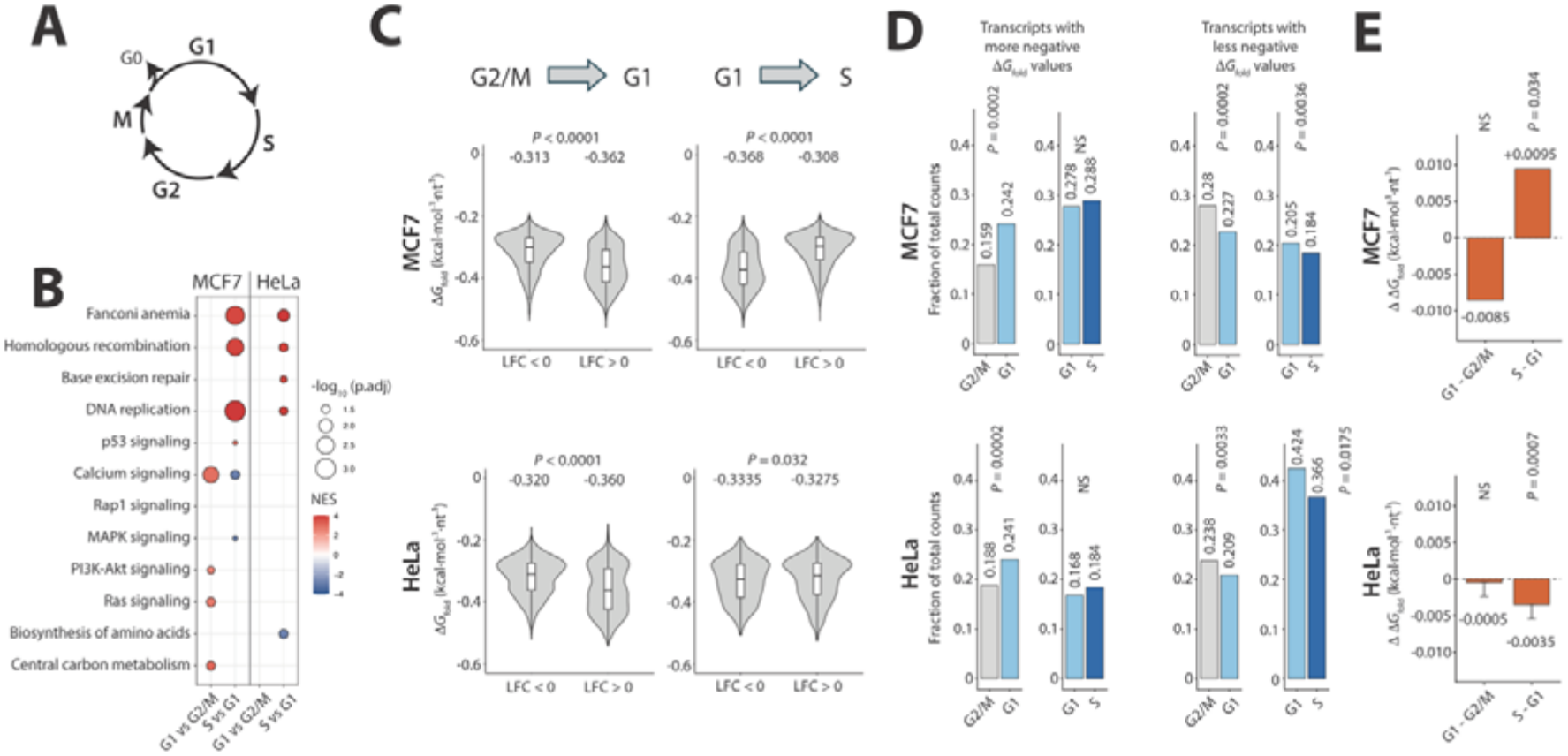
Cell-cycle progression dynamically reweights predicted RNA structural-stability profiles. (**A**) Schematic overview of the human cell cycle, illustrating progression through G1, S, G2 and M phases. (**B**) GSEA of transcriptomes corresponding to selected cell-cycle phase transitions in MCF7 breast cancer and HeLa cervical cancer cells. (**C**) Violin plots showing normalized Δ*G*_fold_ values of transcripts significantly up-or down-regulated across cell-cycle transitions. Values are stratified by log2 fold change (LFC) direction. Wilcoxon rank-sum test. (**D**) Fraction of total normalized read counts contributed by significantly differentially expressed transcripts in the lower or upper Δ*G*_fold_ quartiles across cell-cycle transitions. *P* values from two-sided folding-label permutation tests, BH-adjusted. (**E**) Transcriptome-wide abundance-weighted Δ*G*_fold_ delta across cell-cycle transitions, calculated using TPM values. Values indicate the difference in abundance-weighted Δ*G*_fold_ between the indicated phases. *P* values were calculated by permutation testing and adjusted for multiple comparisons using the Benjamini–Hochberg method.

Within the model, this redistribution of metabolic processes is expected to modulate the balance between internal entropy production and entropy export, such that intracellular configurational entropy may increase or decrease across successive transitions. Accordingly, increased biosynthetic activity is expected to reduce configurational entropy and correspond to more-negative abundance-weighted Δ*G*_fold_ profiles, whereas relative attenuation is expected to increase entropy and correspond to less-negative profiles (Supplementary Information).

To evaluate these model-derived predictions, we systematically screened publicly available RNA-seq datasets in NCBI Gene Expression Omnibus (GEO)^26^ (Methods) and retained studies of two human cancer cell lines: the breast adenocarcinoma line MCF7^27^, showing the expected phase-specific transcriptional changes, and the cervical carcinoma line HeLa^28^, which exhibits atypical S-phase regulation with impaired p53-dependent G1/S control^29–31^. Both cell lines were profiled across G1, S and G2/M phases. Gene- and transcript-level expression data were retrieved through the GEO RNA-seq Experiments Interactive Navigator (GREIN)^32^ platform to ensure uniform reprocessing and minimize technical variability across studies.

To ascertain whether transcriptional signatures reflected the expected metabolic activities and signaling pathways associated with each cell cycle phase, we performed Gene Set Enrichment Analysis (GSEA). At the transcriptional level, both cell lines showed significant S-phase enrichment of DNA replication and repair pathways, including homologous recombination and Fanconi anemia. In contrast, the transition from G2/M to G1 was characterized by the activation of mitogenic signaling and biosynthetic programs, particularly in MCF7 cells (Fig. 2B).

In line with model predictions, the increased biosynthetic activity accompanying the G2/M-to-G1 transition coincided with the upregulation of transcripts with more negative Δ*G*_fold_ values in both cell lines. By contrast, during the G1 to S progression, as DNA replication initiates and broader anabolic programs comparatively attenuate, the Δ*G*_fold_ distribution of upregulated transcripts was biased toward less-negative values, indicating lower predicted secondary-structure stability, predominantly in MCF7 cells (Fig. 2C).

To determine whether these transcript-level shifts were reflected in RNA molecule abundance, we next stratified significantly differentially expressed transcripts into lower and upper Δ*G*_fold_ quartiles and calculated their contribution to total normalized RNA counts. During the G2/M-to-G1 transition, both MCF7 and HeLa cells showed increased RNA-count representation of highly negative Δ*G*_fold_ transcripts, together with reduced representation of less negative Δ*G*_fold_ transcripts, consistent with enrichment of RNAs with greater predicted structural stability in G1 (Fig. 2D). During G1-to-S progression, upper-quartile count fractions decreased within the regulated transcript pools of both cell lines.

At the transcriptome-wide level, the abundance-weighted ΔΔ*G*_fold_ index, defined as the difference in abundance-weighted mean Δ*G*_fold_ between the comparison and reference phases (Methods), was negative during G2/M-to-G1 progression in both cell lines, consistent with the predicted direction, although neither shift was statistically significant. G1-to-S progression produced significant opposing indices: positive in MCF7, also consistent with model predictions, and negative in HeLa, which exhibits atypical S-phase regulation (Fig. 2E).

Together, these findings suggest an association between phase-specific biosynthetic programmes and the reweighting of predicted RNA structural-stability profiles during cell-cycle progression.

### Molecularly targeted anticancer agents reweight the transcriptome toward reduced predicted RNA structural stability

To determine whether analogous RNA-pool reweighting occurs under non-homeostatic conditions that impair cell proliferation, we analyzed GREIN transcriptomic profiles of human cancer cell lines treated with selective inhibitors of proliferative and cell-cycle pathways. From an initial pool of 32 GEO datasets, stringent selection criteria were applied (Methods) yielding 13 datasets spanning 15 human cancer cell lines across 6 histotypes, each treated with one of 13 selective anticancer agents (Fig. 3A and Supplementary Table 3).

**Fig. 3.**
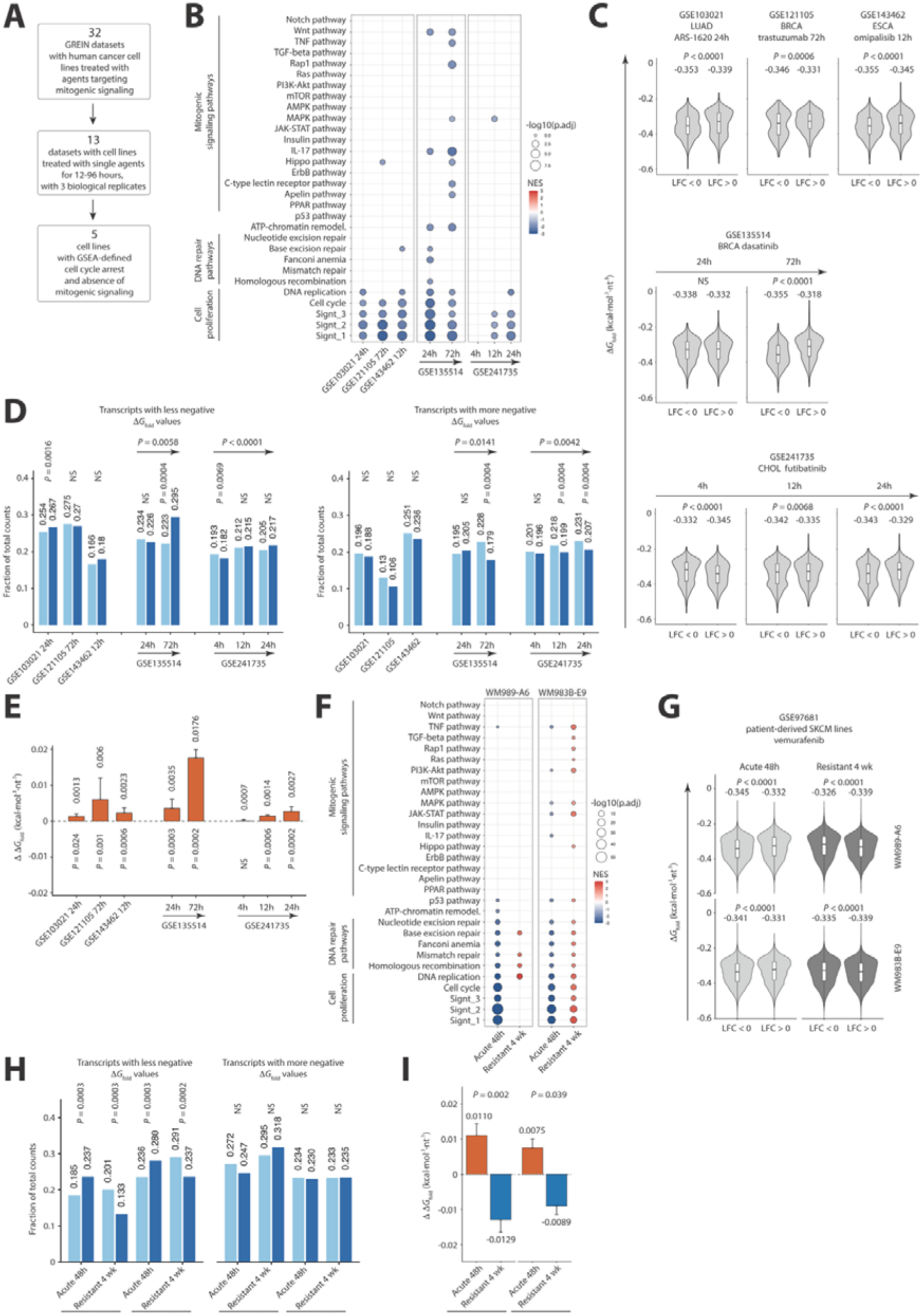
Pharmacologic growth arrest reweights the transcriptome toward reduced predicted RNA structural stability, with an opposing shift during stable drug resistance. (**A**) Selection of GREIN datasets with transcriptional evidence of suppressed proliferation and no residual mitogenic signaling. (**B**) GSEA profiles of the selected treatments; color indicates NES and size indicates −log10(adjusted *P* value). Blank positions indicate nonsignificant enrichment. (**C**) Normalized transcript Δ*G*_fold_ distributions among significantly downregulated and upregulated transcripts. Values above violins are means; *P* values are from two-sided Wilcoxon rank-sum tests. (**D**) Fractions of total normalized counts contributed by upper- and lower-Δ*G*_fold_ quartile transcripts within each significant-DE pool in control and treated samples. *P* values from folding-label permutations for individual comparisons and Welch–Satterthwaite contrasts for temporal changes in treatment–control differences; all two-sided and separately BH-adjusted. (**E**) Transcriptome-wide TPM-weighted ΔΔ*G*_fold_, calculated as treated minus control; *P* values are from permutation tests with Benjamini–Hochberg correction. (**F–I**) Analysis of patient-derived WM989-A6 and WM983B-E9 SKCM lines following acute vemurafenib treatment (48□h) and stable resistance acquired after four weeks under continued drug exposure. (**F**) GSEA profiles for acute versus untreated and stable-resistant versus acute states. (**G**) Corresponding Δ*G*_fold_ distributions among significantly downregulated and upregulated transcripts; light and dark grey denote the acute and stable-resistant comparisons, respectively. *P* values are from two-sided permutation tests. (**H**) Corresponding upper- and lower-quartile normalized-count fractions in the reference and comparison states. *P* values from two-sided folding-label permutation tests, BH-adjusted. (**I**) Transcriptome-wide TPM-weighted ΔΔ*G*_fold_ for the two sequential transitions; orange and blue indicate acute versus untreated and stable resistant versus acute, respectively. *P* values are BH-adjusted one-sided paired *t*-tests across matched founders. Error bars in (E) and (I) indicate propagated SEM. ESCA, esophageal carcinoma; BRCA, breast invasive carcinoma; LUAD, lung adenocarcinoma; CHOL, cholangiocarcinoma; SKCM, skin cutaneous melanoma; LFC, log2 fold change.

Tumors exhibit extensive genetic and phenotypic heterogeneity^33^, a feature mirrored in cancer cell lines. Consequently, even among lines of the same histotype, differences in oncogenic pathways can markedly influence sensitivity to targeted inhibition.

Therefore, to retain only conditions with transcriptional evidence of a cytostatic response, we employed KEGG pathway GSEA, to identify cell lines with down-regulation of multiple proliferation signatures^34–36^ (Supplementary Table 4) and no detectable residual mitogenic signaling. This filtering retained five responsive cell lines from five independent datasets, each displaying a transcriptional profile consistent with growth arrest at one or more time points (Fig. 3B).

Targeted anti-cancer agents typically induce a G1 arrest, suppressing cell-cycle transcriptional programs while activating stress-adaptation pathways. Accordingly, unlike physiological G1 progression, pharmacologic growth arrest is expected to favor transcripts with less negative predicted Δ*G*_fold_ values, consistent with reduced secondary-structure stability.

Across three independent single-time-point datasets, this pattern was highly consistent: upregulated transcripts exhibited significantly less negative Δ*G*_fold_ values in LU65 lung carcinoma cells treated with the KRAS^G12C^ inhibitor ARS-1620 (GSE103021), BT-474 breast carcinoma cells treated with the HER2-targeting antibody trastuzumab (GSE121105), and KYSE-150 esophageal squamous-cell carcinoma cells treated with the dual PI3K–mTOR inhibitor omipalisib (GSE143462) (Fig. 3C).

Analysis of two independent time-course datasets revealed that the shift toward less negative Δ*G*_fold_ values emerged progressively after treatment. In MDA-MB-468 breast cancer cells exposed to the multi-kinase inhibitor dasatinib (GSE135514), the shift was absent at 24 hours but significant by 72 hours. A second time-course dataset, comprising ICC13-7 intrahepatic cholangiocarcinoma cells treated with the FGFR1–4 inhibitor futibatinib (GSE241735), revealed a similar temporal progression with finer resolution: at 4 hours, upregulated transcripts remained enriched for more negative Δ*G*_fold_, whereas a significant shift toward less negative Δ*G*_fold_ values emerged at 12 hours and became more pronounced by 24 hours (Fig. 3C).

We next quantified upper- and lower-Δ*G*_fold_ quartile contributions to total normalized counts within each significantly differentially expressed transcript pool. Significant enrichment of less-negative upper-quartile transcripts occurred with ARS-1620 (GSE103021) and at 72 h with dasatinib (GSE135514), accompanied in the latter by lower-quartile depletion.

Futibatinib (GSE241735) significantly reduced upper-quartile representation at 4 h and lower-quartile representation at 12 and 24 h. In both time-course datasets, treatment–control differences shifted significantly toward upper-quartile enrichment and lower-quartile depletion over time (Fig. 3D). The abundance-weighted ΔΔ*G*_fold_ index for the complete expressed RNA pool was significantly positive in all three single-time-point datasets, indicating net shifts toward less-negative Δ*G*_fold_ values. In time-course models, the index increased from 24 to 72 hours with dasatinib (GSE135514) and became significantly positive by 12 hours with futibatinib (GSE241735), increasing further at 24 hours (Fig. 3E).

To determine whether the shared RNA structural-stability signature reflected similar underlying transcriptional programmes, we compared gene-expression responses across the five models, identifying significant overall differences (*P* = 0.0002). GSEA comparisons also revealed functional heterogeneity (Supplementary Fig. 2A–C).

To further assess the association between RNA-pool reweighting and proliferative state, we revisited the initial pool of 32 datasets and selected treatment conditions that retained transcriptional evidence of mitogenic signaling (Supplementary Fig. 3A).

Under these conditions, two datasets showed significant preferential upregulation of transcripts with more-negative predicted Δ*G*_fold_ values (Supplementary Fig. 3B). Quartile analysis supported this pattern in GSE178755 (Supplementary Fig. 3C). Complete-pool abundance-weighted shifts were negative in three of four conditions but nonsignificant, with their modest magnitude potentially reflecting pharmacological effects on proliferation despite residual mitogenic signaling (Supplementary Fig. 3D). In BT-20 triple-negative breast cancer cells, 24 h of erlotinib treatment (GSE148096) produced a significant shift toward less-negative Δ*G*_fold_ values despite retained mTOR and PPAR signaling. Concurrent suppression of cell-cycle and DNA-replication programs indicated a possible mixed response (Supplementary Fig. 3A), consistent with the reported 20% slowing of population growth^37^.

To formally test whether RNA-pool reweighting tracks proliferative state rather than drug exposure itself, we analysed an additional independent dataset (GSE97681) comprising patient-derived BRAF-V600E melanoma lines WM989-A6 and WM983B-E9 before treatment, after 48_jh of vemurafenib, and after stable resistance had developed following four weeks of continuous drug exposure^38^. Acute treatment downregulated all proliferation signatures without detectable residual mitogenic signaling, whereas stably resistant cultures reactivated proliferative and mitogenic programs while continuing to grow in vemurafenib (Fig. 3F).

In both models, upregulated transcripts had significantly less-negative Δ*G*_fold_ values than downregulated transcripts during acute treatment, but significantly more-negative values during stable resistance (Fig. 3G). Upper-quartile contributions to normalized counts increased significantly during acute treatment and decreased significantly during stable resistance (Fig. 3H).

Notably, the abundance-weighted ΔΔ*G*_fold_ index for the complete RNA pool revealed opposing shifts in both models: positive during acute treatment relative to untreated cells and significantly negative during resistance relative to the acute state (*P* = 0.002 and *P* = 0.039 for WM989-A6 and WM983B-E9, respectively; Fig. 3I). This reversal accompanied renewed proliferation under continued drug exposure, supporting the association between RNA-pool reweighting and proliferative state.

To further assess the generality of the pharmacologically induced RNA-pool reweighting, we analyzed microarray-based perturbation signatures from the iLINCS database^39^ for human cancer cell lines exposed to diverse targeted agents, with gene-expression changes reported as z-scores rather than transcript-abundance measurements.

While this resource enables large-scale comparisons, the design of microarray hybridization inherently limits isoform-level resolution. To mitigate this technical constraint, we selected the Ensembl canonical transcript for each gene as a representative isoform. To assess how closely the canonical-transcript value represented the gene-level isoform ensemble, we compared it with the mean Δ*G*_fold_ across all annotated isoforms. The two measures were strongly correlated (Pearson r = 0.976, *P* < 0.0001; Supplementary Fig. 4), supporting use of canonical-transcript Δ*G*_fold_ as a representative gene-level descriptor in the microarray analysis.

**Figure 4.**
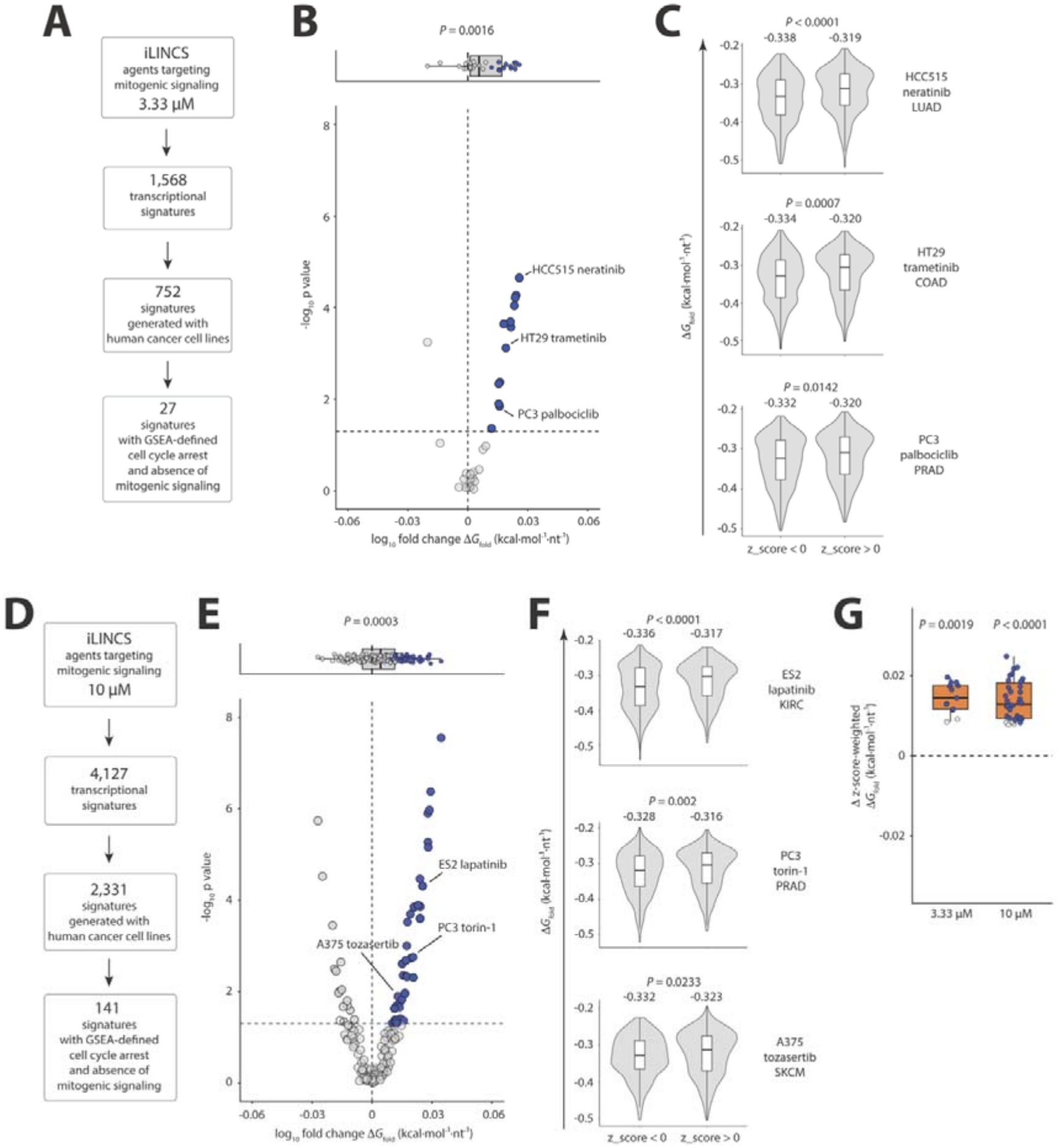
iLINCS signatures reveal transcriptome reweighting toward reduced predicted RNA structural stability under targeted pharmacologic growth arrest. (**A, D**) Flowcharts summarizing the filtering of iLINCS cell line signatures exposed to molecularly targeted agents at 3.33 µM and 10 µM, respectively. (**B, E**) Volcano plots summarizing Δ*G*_fold_-associated transcriptome reweighting across individual cell-line/drug signatures at 3.33 µM and 10 µM, respectively. Blue dots indicate signatures with significant preferential upregulation of genes represented by canonical transcripts with less-negative Δ*G*_fold_ values, corresponding to reduced predicted RNA secondary-structure stability. Dashed horizontal lines denote the statistical significance threshold. Light grey box plots display the distribution of log10 fold changes across the datasets. *P* values from two-sided Wilcoxon signed-rank test. (**C, F**) Violin plots showing the distributions of normalized Δ*G*_fold_ among downregulated (z-score < 0) and upregulated (z-score > 0) transcripts in representative cell-line/drug signatures at 3.33 µM and 10 µM, respectively. Mean normalized Δ*G*_fold_ values are shown above each distribution. *P* values from two-sided Wilcoxon rank-sum tests. **(G)** Z-score-weighted normalized Δ*G*_fold_ perturbation index across iLINCS signatures that significantly up-regulate transcripts with less negative Δ*G*_fold_ values. Positive values indicate less negative normalized Δ*G*_fold_ among positively perturbed genes. Dots represent cell-line/drug conditions; blue dots indicate significant positive shifts. Boxplots summarize each dose group, with two-sided Wilcoxon signed-rank *P* values annotated.

To further validate this approach, we re-analyzed the GREIN RNA-seq datasets, this time restricting Δ*G*_fold_ calculations to canonical transcripts. The resulting predicted RNA structural-stability profiles closely recapitulated those obtained using the complete isoforms set, confirming that the use of canonical transcript Δ*G*_fold_ values preserve the observed patterns of RNA-pool reweighting (Supplementary Fig. 5A-C).

**Fig. 5.**
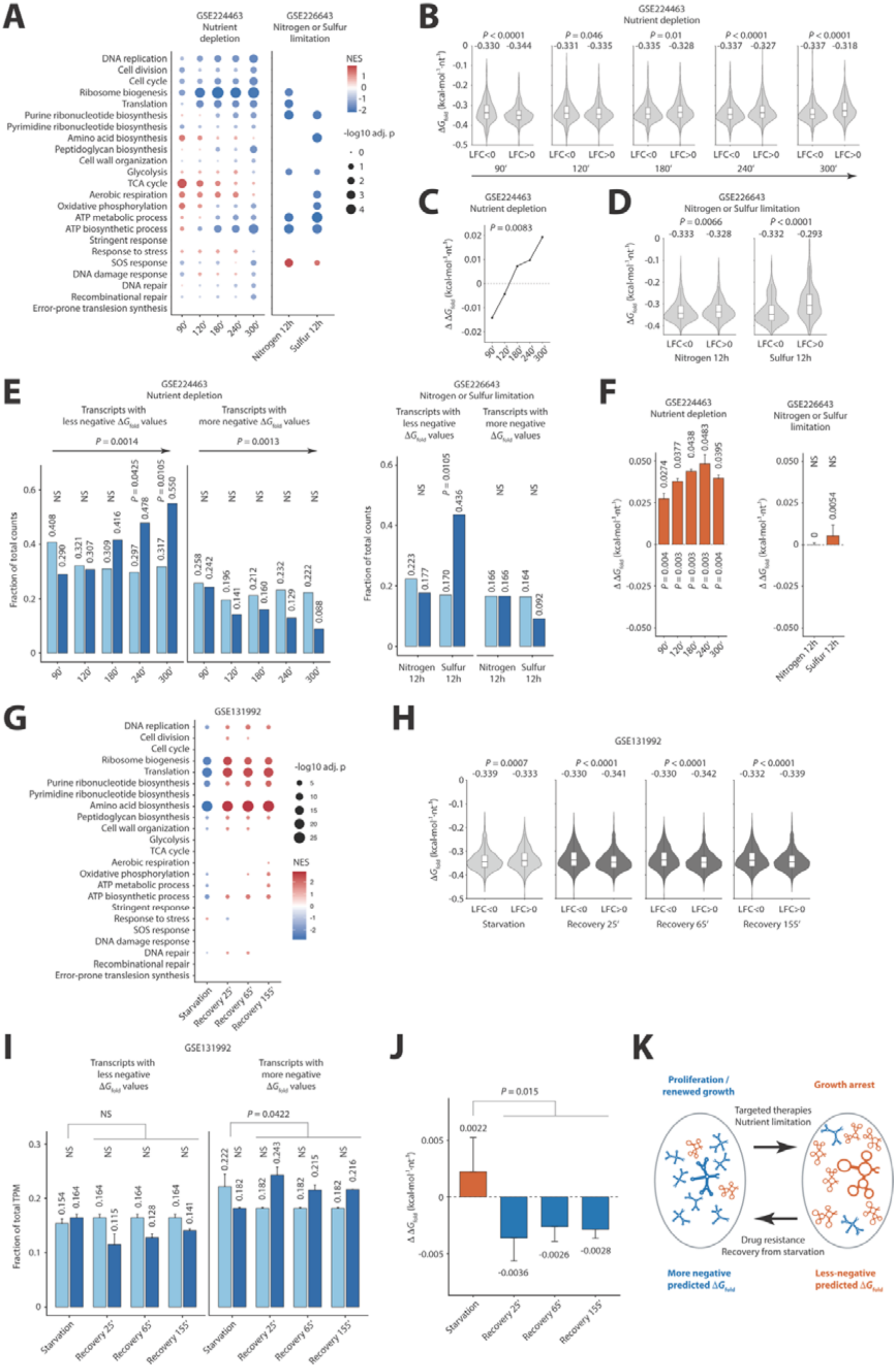
*Escherichia coli* K-12 stationary-phase entry and recovery are accompanied by reversible transcriptome reweighting of predicted RNA structural stability. (**A**) GSEA of selected growth, biosynthetic, metabolic and stress-response pathways during stationary-phase entry. GSE224463 shows 90–300 min relative to the matched 0-min baseline; GSE226643 shows 12-h stationary phase relative to the 3-h early state. (**B**) Normalized Δ*G*_fold_ distributions of FDR-significant transcripts with lower or higher expression across the GSE224463 time course. Mean values are shown above the distributions. Two-sided Wilcoxon rank-sum tests with Benjamini–Hochberg correction. (**C**) Temporal change in the mean normalized Δ*G*_fold_ difference between higher- and lower-expressed significant-DE transcripts from B. One-sided exact Spearman correlation test. (**D**) Normalized Δ*G*_fold_ distributions of FDR-significant transcripts in GSE226643 after nitrogen or sulfur limitation. Two-sided Wilcoxon rank-sum tests with BH correction. (**E**) Fraction of normalized counts contributed by the upper and lower Δ*G*_fold_ quartiles within each comparison-specific significant-DE transcript pool. Light and dark blue indicate the early/baseline and stationary/target states, respectively. *P* values from folding-label permutation tests for individual comparisons and one-sample *t*-tests of replicate slopes for temporal trends in dark-blue fractions; all two-sided and separately BH-adjusted. (**F**) Full-transcriptome RNA-molecule-abundance-weighted Δ*G*_fold_ change. Positive values indicate reweighting toward less negative Δ*G*_fold_. Error bars represent SEM; *P* values from transcriptome-level permutation tests with Benjamini– Hochberg correction. (**G**) GSEA in GSE131992 during established glucose starvation versus pre-starvation and at 25, 65 and 155Dmin of recovery versus established starvation. (**H**) Normalized Δ*G*_fold_ distributions of DESeq2-significant transcripts. Two-sided Wilcoxon rank-sum tests with BH correction. (**I**) Fractions of total TPM contributed by fixed transcriptome-wide upper and lower Δ*G*_fold_ quartiles. Light and dark blue denote reference and target states, respectively. Error bars indicate SEM. Individual comparisons used folding-label permutation tests; mean recovery versus starvation used a Welch–Satterthwaite contrast. Tests were two-sided, with separate BH correction. (**J**) Full-transcriptome abundance-weighted ΔΔ*G*_fold_ during starvation versus pre-starvation (orange) and recovery versus established starvation (blue). Positive and negative values indicate shifts toward less- and more-negative Δ*G*_fold_, respectively; error bars show propagated SEM. **(K)** Schematic of reversible, abundance-driven RNA-pool reweighting during growth arrest and renewed proliferation in human cancer cells and *E. coli*.

Within iLINCS, we selected the longest available exposure (24 hours), providing the closest correspondence to the predominantly 24–72-hour GREIN datasets, and focused on 3.33 and 10 µM, which provided the greatest sample coverage. We then used KEGG pathway GSEA, together with downregulation of proliferation signatures, to identify cell-line/drug conditions consistent with growth arrest and without detectable transcriptional evidence of residual mitogenic signaling (Supplementary Fig. 6A, C).

This multi-step filtering resulted in a final set of 27 transcriptional signatures at 3.33 µM and 141 signatures at 10 µM (Fig. 4A, D), corresponding to 24 and 122 unique cell-line/drug combinations, respectively. The lower drug concentration yielded fewer qualifying signatures, likely reflecting incomplete suppression of mitogenic signaling in a larger proportion of cell lines. At 3.33 µM, the selected conditions comprised 7 cell lines from 5 cancer types treated with 17 targeted agents (Supplementary Fig. 6B), whereas at 10 µM they encompassed 37 cell lines from 12 cancer types treated with 31 targeted agents (Supplementary Fig. 6D).

The iLINCS analysis recapitulated the directional GREIN findings: canonical transcripts representing positively perturbed genes tended to have less-negative predicted Δ*G*_fold_ values than those representing negatively perturbed genes. Across the complete signature sets, positive shifts were observed at both 3.33 and 10 µM (*P* = 0.0016 and *P* = 0.0003, respectively). At 3.33 µM, 22 of 27 signatures (81.5%) shifted in this direction, with 11 (40.7%) individually significant (Fig. 4B, C); at 10 µM, 89 of 141 (63.1%) shifted in the same direction, with 37 (26.2%) individually significant (Fig. 4E, F).

This variability may reflect differences in drug-target expression and cell-line sensitivity, together with the relatively short 24-hour exposure. GREIN time courses showed that the Δ*G*_fold_ shift emerged progressively and strengthened with treatment duration, suggesting that some iLINCS profiles may capture earlier stages of the response.

Among signatures with significant positive shifts, we calculated a ΔΔ*G*_fold_ perturbation index weighted by absolute z-scores, defined as the weighted mean normalized Δ*G*_fold_ of positively perturbed genes minus that of negatively perturbed genes (Methods). The index was positive at both concentrations (*P* = 0.0019 at 3.33 µM and *P* < 0.0001 at 10 µM), indicating that the directional pattern persisted after accounting for perturbation magnitude within this selected subset (Fig. 4G).

As in the GREIN datasets, Spearman-distance, principal-coordinate and variable-pathway analyses revealed heterogeneous pathway responses across cell-line/drug conditions despite the shared structural-stability signature (median pairwise Spearman ρ = 0.290 at 3.33 µM and ρ = 0.241 at 10 µM; Supplementary Fig. 7A–F).

To examine whether the direction of RNA-pool reweighting was associated with mitogenic state, we next analyzed iLINCS signatures from treatment conditions retaining transcriptional evidence of proliferative signaling. In contrast to growth-arrest-associated signatures, these profiles showed a reciprocal tendency at both concentrations, with positive perturbation preferentially observed among genes whose transcripts had more-negative predicted Δ*G*_fold_ values (Supplementary Fig. 8A–F). Although the cohort-level shifts did not reach statistical significance, the z-score-weighted normalized Δ*G*_fold_ index was negative across all qualifying signatures and significantly below zero at 10 µM (*P* = 0.011; Supplementary Fig. 8G), supporting an association between retained mitogenic signaling and reweighting toward greater predicted RNA structural stability.

Together, these data indicate that transcriptome reweighting toward RNAs with less negative Δ*G*_fold_ emerges as a recurrent feature of early responses to targeted anticancer therapies, despite marked line-, drug-, and pathway-level heterogeneity.

### *Escherichia coli* stationary-phase entry recapitulates the RNA thermodynamic signature of growth arrest

The consistency of this directional shift across diverse human cancer cell models raises the question of whether it reflects a fundamental physical constraint. If so, an analogous signature should emerge in phylogenetically distant organisms despite profound differences in their gene-regulatory networks and metabolic architectures. We therefore systematically screened GEO^26^ and identified three eligible RNA-seq datasets of *Escherichia coli* K-12 capturing ppGpp/RpoS-associated stress-adaptive growth arrest^40,41^ (Methods). Two comprised nutrient depletion over 0–300_jmin (GSE224463) and 12-h nitrogen or sulfur limitation (GSE226643).

Δ*G*_fold_ values were computed for 4,570 *E. coli* genes using the ViennaRNA workflow (Methods and Supplementary Table 7).

Across the two datasets, GSEA identified repression of growth-related, biosynthetic and bioenergetic programs, with the affected pathways varying by time point and nutrient limitation (Fig. 5A).

In the GSE224463 nutrient-depletion time course, the association between differential expression and predicted RNA stability changed over time. Consistent with the temporal GSEA profile, at 90 and 120 min, significantly upregulated transcripts had more negative Δ*G*_fold_ values than significantly downregulated transcripts. As repression broadened from 180 min onward to encompass translation, anabolic pathways and ATP biosynthesis, this relationship reversed, and upregulated transcripts became progressively enriched for less-negative values (Fig. 5A, B).

The mean Δ*G*_fold_ difference between significantly upregulated and downregulated transcripts showed a strong positive association with time (Spearman ρ = 1, *P* = 0.0083; Fig. 5C).

Within the comparison-specific regulated RNA pools, upper-quartile count fractions increased and lower-quartile fractions decreased overall across 90–300 min (*P* = 0.0014 and 0.0013, respectively), with upper-quartile enrichment relative to baseline reaching significance at the final two time points, 240 and 300 min (Fig. 5E).

At the full-transcriptome level, the abundance-weighted ΔΔ*G*_fold_ index was significantly positive at all five time points, indicating early net RNA-pool reweighting before reversal of the Δ*G*_fold_ pattern among differentially expressed transcripts (Fig. 5F).

GSE226643 reproduced this directional shift. Under nitrogen limitation, transcripts upregulated at 12 h had modestly but significantly less negative Δ*G*_fold_ values than downregulated transcripts (*P* = 0.0066), with a more pronounced separation under sulfur limitation (*P* < 0.0001; Fig. 5D). Among the corresponding quartile comparisons, the upper-quartile enrichment under sulfur limitation was significant (*P* = 0.0105; Fig. 5E). The corresponding full-transcriptome abundance-weighted ΔΔ*G*_fold_ indeces were directionally concordant but not significant (Fig. 5F).

To formally test whether this shift reverses during recovery from growth arrest, we analyzed the third eligible dataset GSE131992^42^. Established glucose starvation suppressed growth-related programs and reweighted both regulated and complete RNA pools toward less-negative Δ*G*_fold_, whereas the three sampled recovery time points, early (25 min), intermediate (65 min) and late (155 min) after glucose replenishment, showed reactivation of growth programs and reversal of these shifts (Fig. 5G– J).

Mean recovery fractions showed a nonsignificant upper-quartile reduction and significant lower-quartile enrichment (*P* = 0.0422; Fig. 5I). The abundance-weighted ΔΔ*G*_fold_ index was positive during starvation and negative at all three recovery time points, with the mean recovery index significantly lower than the starvation index (*P* = 0.015; Fig. 5J).

Importantly, across all three *E. Coli* datasets, the four nutrient-limited growth-arrest conditions, the shared RNA structural-stability signature emerged despite significant overall differences in pathway-level transcriptional responses (*P* = 0.0065; Supplementary Fig. 9A–C).

Together, these findings show that pharmacologically induced cell-cycle arrest in human cells and nutrient-limited stationary-phase entry in *Escherichia coli* converge on a common directional reweighting of transcript abundance toward RNAs with less-negative predicted Δ*G*_fold_ values. The reversal of this shift upon renewed proliferation in both systems supports its interpretation as a conserved, reversible signature of growth state across distinct stressors (Fig. 5K).

## Discussion

From a physical perspective, cells are open systems whose far-from-equilibrium organization is sustained by continuous free energy input and entropy export to the environment. Thermodynamic principles underpin fundamental molecular mechanisms including protein folding^43^, enzyme kinetics^44^, molecular motor activity^45^, transcription^46^ and translation^47^. Non-equilibrium principles have also been applied to regulatory networks controlling cell fate, including the cyclin-dependent kinase network during cell cycle progression^48^ and the rhythmic synthesis of the tumor suppressor p53^49^.

Beyond these molecular mechanisms, thermodynamic constraints may be deeply embedded within higher-order cellular processes. These include cell cycle progression and homeostatic responses, which entail complex metabolic reprogramming, signaling rewiring and transcriptional changes.

Among these, the transcriptional response to selective pressures is particularly relevant: it constitutes a central mechanism mediating survival and adaptation while integrating upstream signaling, metabolic state and environmental inputs, making the transcriptome an informative system in which to investigate whether thermodynamic constraints contribute to cellular adaptation over time.

The central role of RNA is further underscored by its evolutionary history^50,51^. In primitive protocells, RNA served both as a carrier of hereditary information and as a catalyst for chemical reactions. Given this centrality, early RNA molecules would have been subject to selection for structures capable of supporting information storage and catalytic function, while physical constraints associated with non-equilibrium conditions may also have shaped the range of accessible structural configurations. From this perspective, the transcriptome emerges as a dynamic interface through which thermodynamic principles could be integrated into core regulatory programs, raising the possibility that the collective biophysical profile of the expressed RNA pool could be reshaped by transcriptional changes.

To examine this possibility, we used predicted Δ*G*_fold_ as a sequence-derived descriptor of RNA secondary-structure stability and interpreted its abundance-weighted profile as a transcriptome-level projection of intracellular configurational change. Within our framework, the predicted Δ*G*_fold_ of each transcript remains a fixed property across conditions, while changes in relative abundance reweight the expressed RNA pool across this distribution of folding stabilities.

Cellular-state transitions involve coordinated reorganization of proliferative and metabolic programs that may alter the balance between internal entropy production and outward entropy flow (Supplementary Information). Within the model, intracellular configurational entropy can increase transiently if entropy export declines more strongly than internal entropy production. Under the proposed projection, this increase would be accompanied by transcriptome reweighting toward less-negative predicted Δ*G*_fold_ values.

Cell-cycle progression provides a physiological reference for this organization. In MCF7 cells, biosynthetically active phases were biased toward transcripts with more negative Δ*G*_fold_ values, whereas transitions involving reduced anabolic activity favoured less negative profiles. These phase- and cell-context-dependent shifts are compatible with reversible reweighting of the RNA pool as anabolic demands change.

This physiological remodelling makes the recurrent directional shift observed during pharmacologically induced growth arrest particularly informative. Across multiple cancer histotypes and molecularly targeted agents, the expressed RNA pool shifted toward less-negative predicted folding free energies despite marked heterogeneity in the pathways affected. Conditions retaining residual mitogenic signaling tended to preserve or regain enrichment for transcripts with more negative Δ*G*_fold_ values, supporting an association with proliferative state rather than drug exposure alone.

The same directional shift was also observed during nutrient-limited *E. coli* stationary-phase entry, extending the observation across human and bacterial systems with fundamentally different transcriptional landscapes and regulatory networks. Opposing shifts during acquired resistance under continued drug exposure and bacterial recovery after glucose replenishment further support an association between RNA-pool reweighting and growth state.

Together, these observations suggest that heterogeneous transcriptional responses elicited by distinct selective pressures can converge on a common ensemble-level biophysical signature as cells transition toward low-growth states.

The parallel extended beyond directionality to temporal progression. In both drug-treated human cells and nutrient-depleted *E. coli*, the shift toward less-negative abundance-weighted Δ*G*_fold_ became progressively more pronounced as the cytostatic or stationary state developed.

This temporal concordance suggests that abundance-weighted Δ*G*_fold_ may track the transcriptome reorganization accompanying the progression from reduced growth to complete arrest and represent a candidate thermodynamic pseudo-time for this trajectory.

These findings motivate an experimentally testable hypothesis. Drug-induced reweighting of the expressed RNA pool toward transcripts with less-stable predicted secondary structures may sensitize malignant cells to mild hyperthermia, given the intrinsic temperature dependence of RNA folding^52–54^. Combining molecularly targeted agents with controlled thermal pulses in human cancer cell lines may constrain this remodeling, thereby limiting adaptive responses, particularly when heat exposure is timed to coincide with peak transcriptional reprogramming, which our data suggest occurs 24–72 hours after treatment initiation.

The potential scope of this thermal sensitization will depend on how the abundance-weighted Δ*G*_fold_ shift is distributed across the treated population. Given the substantial genetic and phenotypic heterogeneity that can persist even within established human cancer cell lines^55^, a related question for future work is whether the abundance-weighted Δ*G*_fold_ shifts observed in bulk populations occur broadly across individual cells or are driven by discrete subpopulations following distinct transcriptome-state trajectories. Single-cell RNA sequencing with sufficient depth and isoform resolution could distinguish between these possibilities.

An important extension would be to determine whether analogous reweighting occurs among non-coding RNAs, whose regulatory functions are often structure dependent^56^.

Beyond the transcriptome, the relationship between cellular state and ensemble-level biophysical organization may extend to protein conformational landscapes^57^ and lipid-bilayer organization^58^. Establishing whether these molecular systems undergo coordinated remodelling during adaptive responses^59^ including the emergence of drug-tolerant persister states^60^ or entry into dormancy^61^ will define the broader applicability of this thermodynamic framework.

Collectively, these findings offer a new lens on the dynamics of cell cycle progression and homeostatic responses, shifting the focus beyond canonical (epi)genetic and molecular mechanisms toward deeper physical constraints that may uncover previously unrecognized therapeutic vulnerabilities.

## Materials and Methods

### Computational prediction of RNA folding stability

RNA secondary-structure predictions were performed across the human transcriptome using transcript sequences from GENCODE v47. For each transcript, the minimum folding free energy (Δ*G*_fold_) was computed in kcal·mol^−1^ with the RNAfold program in ViennaRNA v2.6.4, run in non-graphical mode with default energy parameters at 37°C. To assess robustness across software implementations, the same transcript sequences were also analysed using the Fold program in RNAstructure v6.5 with default RNA energy parameters at 37°C. Both programs use dynamic programming to predict minimum-free-energy secondary structures under nearest-neighbour thermodynamic models, with differences in implementation and energy scoring. For both programs, predicted folding free energies were normalized by transcript length and expressed as length-normalized Δ*G*_fold_ in kcal·mol^−1^·nt^−1^.

### PARIS2 analysis of HEK293T mRNA duplexes

Publicly available oligo(dT)-enriched HEK293T PARIS2 data generated after amotosalen-mediated *in vivo* RNA crosslinking were used to evaluate whether experimentally captured mRNA duplexes were supported by transcript-level ViennaRNA secondary-structure predictions. CRSSANT^62^-derived BEDPE duplex groups representing proximity-ligated, gapped RNA fragments were used as the structural readout of RNA duplex formation. The analysis was restricted to protein-coding intramolecular mRNA duplex groups with CRSSANT support ≥2. CRSSANT BEDPE duplex intervals were filtered for same-gene, same-strand intramolecular interactions and mapped to GENCODE v47 exon models to convert genomic duplex arms into transcript-coordinate intervals. Each duplex arm was required to overlap the corresponding transcript exon model by at least 5 nucleotides. Only duplexes for which both arms mapped to the same transcript were retained. ViennaRNA dot-bracket structures were converted into transcript-level base-pair tables, and each mapped PARIS2 duplex was scored as supported when at least one predicted ViennaRNA base pair connected the two experimentally defined duplex arms. Support was assessed using exact coordinate matching and relaxed windows of ±5, ±10 and ±15 nucleotides. As a matched null model, each experimental duplex was randomized within its own transcript by repositioning the two duplex arms while preserving the original arm lengths and the distance separating them. ViennaRNA support for these geometry-matched random duplexes was quantified using the same overlap criteria over 1,000 randomizations. For each matching window, the observed support was compared with the resulting null distribution using a one-sided empirical *P* value, and the 2.5th and 97.5th percentiles of the null distribution were calculated as a 95% null interval.

### icSHAPE-based analysis of *in vivo* RNA accessibility

Publicly available transcriptome-wide *in vivo* icSHAPE datasets from H9, HeLa, HepG2 and K562 cells, together with an icSHAPE-MaP dataset from HEK293T cells, were used as independent experimental measures of nucleotide-level RNA accessibility. Raw strand-specific genomic signal tracks were processed as BED-like one-nucleotide intervals, in which each row represented a genomic position, strand, and icSHAPE reactivity score. These genomic scores were mapped to transcript coordinates using GENCODE v47 exon annotations with strand-aware exon ordering. Genomic positions overlapping more than one strand-compatible transcript model were assigned to each compatible transcript and converted to the corresponding transcript coordinates. In parallel, ViennaRNA dot-bracket structures were expanded into per-nucleotide transcript tables, and each position was classified as paired or unpaired according to the predicted secondary structure. Transcript-mapped icSHAPE scores were then joined to ViennaRNA structural states by matched transcript ID and transcript coordinate. icSHAPE and icSHAPE-MaP reactivity values were analyzed as continuous nucleotide-level accessibility measurements and were not dichotomized into experimental paired or unpaired calls. Only transcripts containing at least 10 icSHAPE-covered positions predicted to be paired and at least 10 icSHAPE-covered positions predicted to be unpaired were retained for paired-versus-unpaired interpretation. For each retained transcript, mean and median icSHAPE reactivity values were calculated separately for ViennaRNA-predicted paired and unpaired positions. Within each transcript, reactivity distributions at predicted paired and unpaired positions were compared using two-sided Wilcoxon rank-sum tests, with Benjamini–Hochberg correction across tested transcripts within each cell line. At the cell-line level, transcript-specific unpaired-minus-paired reactivity differences were tested against zero using one-sided Wilcoxon signed-rank tests. Concordance was defined as higher icSHAPE reactivity at ViennaRNA-predicted unpaired positions than at predicted paired positions, consistent with the expected greater chemical accessibility of unpaired RNA nucleotides.

### Matched shuffled-sequence controls for RNA folding stability

To control for sequence-composition effects on predicted RNA folding stability, matched shuffled-sequence controls were generated for transcript-level native-versus-shuffled comparisons. Full transcript sequences were shuffled while preserving dinucleotide composition, and each shuffled replicate was folded with RNAfold using the same ViennaRNA settings applied to the native sequences. For each transcript, shuffled-control summaries were calculated from matched shuffled replicates, with a standard design of 20 shuffled sequences per native transcript. Native-minus-shuffled folding values were computed on a transcript-matched basis as Δ*G*_fold_ native minus the mean Δ*G*_fold_ shuffled, and length-normalized Δ*G*_fold_ native minus the mean length-normalized Δ*G*_fold_ shuffled. Negative native-minus-shuffled values therefore indicate greater predicted stability of the native transcript relative to matched dinucleotide-preserving shuffled controls, whereas positive values indicate lower native stability relative to the matched shuffled expectation.

### RNA biotype and pathway-level analyses of structural stability

To prevent genes with many annotated isoforms from disproportionately weighting the biotype-level stability distributions, analyses were restricted to one canonical or representative transcript per gene. Normalized Δ*G*_fold_ was compared among protein-coding transcripts, tRNAs, and rRNAs using Wilcoxon rank-sum tests with Benjamini–Hochberg correction. Native-minus-shuffled normalized Δ*G*_fold_ values were compared with zero within each biotype and across biotypes using Wilcoxon tests. For pathway analysis, one representative protein-coding transcript per gene was ranked by native-minus-shuffled normalized Δ*G*_fold_, and genes in the most- and least-stable 10% were subjected to over-representation analysis against Reactome and KEGG pathway annotations using clusterProfiler R package, with Benjamini–Hochberg correction.

### RNA secondary structure visualizations

For selected transcripts, secondary structure predictions were performed using RNAfold (ViennaRNA package, v2.6.4) via the Web Services portal (http://rna.tbi.univie.ac.at). Predictions were computed using default thermodynamic parameters, with the partition function calculation enabled and the no lonely pairs constraint applied to avoid isolated base pairs. Centroid secondary structures—defined as the structure minimizing the Boltzmann-weighted average base-pair distance to all structures in the ensemble—were extracted from the resulting partition function ensemble. Structures were generated for length-matched protein-coding transcripts at the upper and lower extremes of the normalized Δ*G*_fold_ distribution and rendered using the default visualization settings of the ViennaRNA Web interface.

### Cell cycle phase-specific analysis of RNA structural stability and pathway activity

The NCBI Gene Expression Omnibus (GEO)^26^ was systematically searched up to 31 July 2025 using NCBI E-utilities and overlapping queries targeting human bulk RNA-seq, cell-cycle phases and experimental phase resolution. Deduplicated GSE records were manually screened. Studies were restricted to peer-reviewed analyses of established human cancer cell lines with phase-resolved sampling. Primary or patient-derived material and experimentally introduced perturbations of endogenous gene expression or genome editing were excluded, whereas reporter constructs used solely for phase identification or sorting were permitted. De novo differential-expression analysis required at least two independent biological replicates per phase. Transcript-level expression data of synchronized human breast (MCF-7; GEO GSE94479) and cervical (HeLa; GEO GSE104736) cancer cells were retrieved from GEO and Sequence Read Archive (SRA)^63^ via the GEO RNA-seq Experiments Interactive Navigator (GREIN)^32^ platform (https://www.ilincs.org/apps/grein/). Samples were grouped into G1, S and G2/M phases as annotated by the original authors, and phase-to-phase differential expression was computed with DESeq2 (v1.40). Length-normalized RNA-folding free energies (Δ*G*_fold_/nt) were merged onto each gene via transcript IDs. Phase-specific pathway activity was assessed by Gene Set Enrichment Analysis (clusterProfiler v4.8) against KEGG, using ranked Wald statistics and Benjamini–Hochberg correction; normalized enrichment scores (NES) and –log_10_ FDR values were visualized as bubble plots. Δ*G*_fold_ distributions of significantly up- and down-regulated transcripts were compared using two-sided Wilcoxon rank-sum tests (Fig. 2C). For quartile analysis (Fig. 2D), lower and upper Δ*G*_fold_ quartiles were defined within each comparison’s significant-DE transcript pool (adjusted *P* < 0.05). Quartile fractions were calculated from phase-averaged DESeq2-normalized counts relative to total counts in the same pool. Phase differences were assessed using 10,000 folding-label permutations, keeping abundance profiles fixed and applying the same label reassignment to both phases. Two-sided empirical *P* values used the plus-one correction and Benjamini–Hochberg adjustment across eight tests.

### Transcriptome-wide abundance-weighted Δ(Δ*G)*_fold_ index analysis

Transcript-level TPM values were used to calculate the abundance-weighted mean predicted folding free energy for each sample *s*:

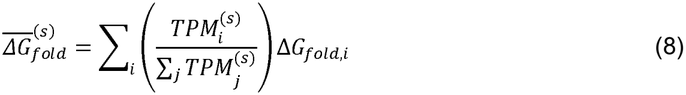

Here, *i* identifies each retained transcript, and the denominator sums TPM values across all retained transcripts, indexed by *j*, in sample (s). ΔG_*fold,i*_ denotes the fixed, length-normalized predicted folding free energy of transcript (s). The abundance-weighted Δ(Δ*G)*_*fold*_ index was defined as the mean abundance-weighted Δ*G*_fold_ across biological replicates of the comparison condition minus that of the reference condition, with propagated standard errors:

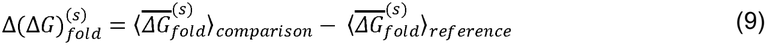

Angle brackets denote means across biological replicates. Positive and negative indices indicate shifts toward less-negative and more-negative Δ*G*_fold_ profiles, respectively. Statistical significance was assessed using 5,000 two-sided permutations in which Δ*G*_fold_ values were reassigned across transcripts while the observed TPM-derived contrast weights were held fixed. Empirical *P* values were adjusted using the Benjamini–Hochberg method.

### Transcriptomic responses of human cancer cells exposed to molecularly targeted agents

Transcript- and gene-level raw and normalized counts were retrieved from the GREIN platform^32^ by querying for human cancer cell lines treated with single small-molecule inhibitors targeting proliferative or cell cycle–associated pathways.

An initial pool of 32 datasets was refined using stringent criteria requiring: i) treatment with a single pharmacologic agent; ii) confirmed presence of the drug’s molecular target(s) in each line; iii) exposure times between 12 and 96 hours; iv) exclusion of cell lines engineered with shRNA or oncogene expression vectors; v) availability of at least two biological replicates per condition; and vi) publication in a peer-reviewed journal. Applying these criteria, we selected 13 datasets encompassing 15 human cancer cell lines across six histotypes, treated with 13 distinct anticancer agents (Table S3).

Sample annotations and metadata were obtained using the GEOquery R package. Raw counts were filtered to retain transcripts with a minimum of 3 reads across all samples. Differential expression analysis was performed using DESeq2 pipeline (v1.40).

To identify treatment conditions associated with cell-cycle arrest, ranked gene lists were analyzed by GSEA using clusterProfiler v4.8. Profiles showing significant negative enrichment of the KEGG Cell Cycle pathway and no significant positive enrichment of selected mitogenic or proliferative KEGG pathways were retained (Benjamini–Hochberg FDR < 0.05). The complete GSE241735 futibatinib time course was retained independently to assess the temporal response. Transcriptional response heterogeneity was tested by partial redundancy analysis of the dataset-by-treatment interaction, controlling for main effects, with 10,000 within-dataset residual permutations. Time courses were represented by their latest time points.

For quartile analysis (Fig. 3D, H), lower and upper Δ*G*_fold_ quartiles were defined among significantly differentially expressed transcripts (adjusted *P* < 0.05). Condition-averaged DESeq2-normalized counts were summed within each quartile and divided by the total across the same retained transcript pool. Between-state differences were assessed using 10,000 two-sided folding-label permutations, keeping abundance profiles fixed and applying the same label reassignment to both states. Empirical *P* values used the plus-one correction and Benjamini–Hochberg adjustment.

### Analysis of RNA structural stability in pharmacologically perturbed cancer transcriptomes from iLINCS

Gene-level expression data of human cancer cell lines treated with molecularly targeted agents were retrieved from the iLINCS database^39^ (https://www.ilincs.org). Analyses were restricted to signatures obtained after 24 hours of treatment at concentrations of 3.33 µM or 10 µM. The same curation criteria were applied to both concentration subsets: normal or composite (“XC.”) cultures and poorly annotated tumor lines were excluded, and signature identifiers were cross validated against official iLINCS metadata.

For the 3.33 µM subset, 1,998 gene-expression signatures representing 39 cell lines, 50 agents, and 911 unique cell line–drug combinations were retrieved. Restriction to the targeted compounds considered in this study yielded 1,568 signatures from 37 cell lines treated with 39 agents, corresponding to 730 unique cell line–drug combinations (Supplementary Table 5). Application of the common curation criteria reduced this collection to 752 cancer-cell-line signatures. The resulting dataset comprised 20 cancer cell lines spanning 12 primary disease categories and 497 unique cell line–drug combinations involving 39 molecularly targeted anticancer agents. Subsequent application of the growth-arrest criteria retained 27 signatures representing 7 cancer cell lines spanning 5 primary disease categories and 4 annotated subtype categories, 17 targeted agents, and 24 unique cell line– drug combinations.

For the 10 µM subset, 7,113 gene-expression signatures representing 157 cell lines, 54 agents, and 3,134 unique cell line–drug combinations were retrieved. Restriction to the targeted compounds considered in this study yielded 4,127 signatures from 151 cell lines treated with 38 agents, corresponding to 2,465 unique cell line–drug combinations (Supplementary Table 5). Application of the common curation criteria reduced this collection to 2,331 cancer-cell-line signatures. The resulting dataset comprised 98 cancer cell lines spanning 22 primary disease categories and 1,536 unique cell line–drug combinations involving 38 molecularly targeted anticancer agents (Supplementary Table 6). Subsequent application of the growth-arrest criteria retained 141 signatures representing 37 cancer cell lines spanning 12 primary disease categories and 10 annotated subtype categories, 31 targeted agents, and 122 unique cell line–drug combinations.

Signatures were computed using the Characteristic Direction (CD) method implemented in an automated Python workflow following the LINCS protocol^64^. Level 3 L1000 data—comprising quantile-normalized and log_2_-transformed gene-expression values—were processed in Python v3.11. For each perturbation, treatment instances were identified according to cell line, compound, dose, and time point, whereas the corresponding control instances were selected from the same experimental batch. Differential-expression vectors were then computed using the CD algorithm from the maayanlab-bioinformatics package. Gene Set Enrichment Analysis was performed using the R package clusterProfiler v4.8 against a prespecified panel of KEGG pathways related to cell-cycle progression and mitogenic signaling, together with three independent proliferation gene sets. Samples were eligible for further analysis when significant enrichment results were available for the KEGG Cell Cycle pathway (hsa04110) and all three proliferation gene sets. Signatures exhibiting negative normalized enrichment scores for all four gene sets were retained, after which samples showing significant positive enrichment of any of 32 prespecified pathways associated with mitogenic signaling or proliferation were excluded. Enrichment was considered significant at a Benjamini–Hochberg FDR below 0.05.

To examine complementary conditions with transcriptional evidence of retained proliferative or mitogenic signaling (Supplementary Fig. 8), eligible signatures were required to have a positive NES for the KEGG Cell Cycle pathway, at least one of the three proliferation gene sets, or one of six major mitogenic pathways: mTOR, Ras, MAPK, PI3K–Akt, insulin or ErbB signaling. Signatures with positive NES for ECM–receptor interaction, calcium signaling, VEGF signaling or apoptosis were excluded. This selection yielded 7 signatures at 3.33 µM and 27 signatures at 10 µM, which were analyzed using the same Δ*G*_fold_ framework described below.

Each gene feature was annotated with the predicted minimum free energy (Δ*G*_fold_, kcal mol^−1^) and sequence length of its corresponding transcript using precomputed values generated with the ViennaRNA package, as described above. Structural comparisons were restricted to genes with a CD-derived nominal *P* value below 0.05. Genes with positive and negative CD coefficients were classified as upregulated and downregulated, respectively. Within each signature, the transcript-length-normalized Δ*G*_fold_ distributions of the two groups were compared using a two-sided Wilcoxon rank-sum test. The magnitude and direction of the structural shift were summarized as log_10_(mean normalized Δ*G*_fold_ of downregulated genes/mean normalized Δ*G*_fold_ of upregulated genes). Positive values therefore indicated that upregulated genes had higher—or less negative—normalized Δ*G*_fold_ values than downregulated genes.

As a complementary effect-size analysis, normalized Δ*G*_fold_ values were weighted by the absolute magnitude of the CD-derived score separately for genes with positive and negative scores. The z-score-weighted Δ*G*_fold_ shift was calculated as the weighted mean for genes with positive scores minus the corresponding weighted mean for genes with negative scores. Significance for each signature was assessed using 1,000 permutations in which normalized Δ*G*_fold_ values were randomly reassigned among genes while the observed CD-derived scores were held fixed; two-sided empirical *P* values were calculated from the proportion of permuted absolute shifts at least as large as the observed absolute shift. For the growth-arrest analysis, this approach was restricted to significant positive-shift signatures, comprising 11 signatures at 3.33 µM and 37 signatures at 10 µM, and group-level shifts were evaluated using two-sided one-sample Wilcoxon signed-rank tests. For the complementary analysis of signatures retaining proliferative or mitogenic activity (Supplementary Fig. 8G), the weighted analysis was restricted to significant more-negative-shift signatures, comprising 3 signatures at 3.33 µM and 7 signatures at 10 µM. Consistent with the directional hypothesis, these group-level indices were evaluated against zero using one-sided Wilcoxon signed-rank tests with the alternative hypothesis of a negative shift.

### Analysis of *Escherichia coli* stationary-phase transcriptomes

GEO was searched through 31 July 2025 using NCBI E-utilities for *E. coli* RNA-seq studies of nutrient limitation, stationary-phase entry and recovery. Peer-reviewed studies of author-designated wild-type K-12 backgrounds with accessible counts or reads, suitable untreated controls and at least two biological replicates per condition were screened manually.

GSE224463 comprised three independent experiments using author-designated wild-type samples (JH01), sampled at 90, 120, 180, 240 and 300 min against matched 0-min baselines. GSE226643 compared 12-h nitrogen-or sulfur-limited cultures with 3-h controls (two biological replicates per condition). ViennaRNA predictions for the MG1655 reference were matched by locus tag. Induced and repressed Δ*G*_fold_ distributions were compared using two-sided Wilcoxon rank-sum tests with BH correction.

Quartiles were defined within each regulated set; summed mean DESeq2-normalised counts were divided by the corresponding pool total. Differences between states were tested using 10,000 two-sided folding-label permutations, holding expression fixed (BH correction across 14 tests). GSE224463 trends used one linear slope per experiment for target-state fractions across 90–300 min, tested against zero using two-sided one-sample *t*-tests (BH correction across two quartiles). The temporal trend in the induced-minus-repressed mean Δ*G*_fold_ difference used one-sided exact Spearman correlation.

For full-transcriptome indices, raw counts were divided by gene length and normalised within samples to relative abundances *p*□. The index was Σ*p*□*x*□, where *x*□ denotes length-normalised predicted Δ*G*_fold_. Index differences were assessed by 5,000 two-sided descriptor-label permutations, BH-adjusted within each dataset, with confidence intervals from 5,000 gene-contribution bootstrap resamples. Replicate-level sensitivity analyses used two-sided paired *t*-tests and paired Wilcoxon tests for GSE224463 and Welch’s *t*-tests for GSE226643.

GSE131992 compared established starvation with pre-starvation and recovery at 25, 65 and 155 min with established starvation. Predictions were recalculated for 4,498 deposited gene intervals. Distribution comparisons excluded rRNAs and required DESeq2 BH-adjusted *P* < 0.05, |log_2_ fold change| ≥ 1 and mean TPM ≥ 1; two-sided Wilcoxon tests were BH-adjusted. Fixed transcriptome-wide quartile fractions and abundance-weighted indices used sample-level relative TPM. Individual quartile comparisons used 10,000 two-sided folding-label permutations (BH correction across eight tests). Mean recovery versus starvation was evaluated using two-sided Welch–Satterthwaite contrast of sample-level fractions, equally weighting the three recovery means and assuming independent sampling errors (BH correction across two quartiles). Index differences were summarised with propagated SEM.

GSEA used clusterProfiler and curated K-12 GO annotations, ranking genes by DESeq2 log_2_ fold change for stationary-phase datasets and Wald statistic for recovery, with BH correction. Cross-dataset functional comparisons used Wald-ranked GO/KEGG enrichment at 300 min, nitrogen limitation and sulfur limitation. Complete NES profiles were compared by Spearman correlation; 5,000 conditional randomisations preserving responsive-term numbers and directions assessed the shared response.

### Software and computational environment

Analyses used R (v4.4.2), with GEOquery, biomaRt, dplyr, tidyr, DESeq2, clusterProfiler, ReactomePA and ggplot2. LINCS L1000 data were additionally processed in Python using cmapPy for data extraction and maayanlab_bioinformatics for Characteristic Direction analysis, supported by pandas, NumPy and SciPy. RNA-folding calculations were performed with ViennaRNA v2.6.4 on the UNSW Katana high-performance computing cluster.

## Acknowledgements

We thank Dr. Mark Pinese, Prof. John Mattick and Prof. Huai Yong Zhu for their critical review of the manuscript, and Dr. Erin Heyer for assistance with bioinformatic analyses.

## Author contributions

Study conceptualization: A.C. Stochastic thermodynamic modelling: R.Z. and A.C. Bioinformatic analyses: A.C., J.R., S.P., A.M.R., A.X., D.M.T., I.D. Manuscript writing: A.C.

## Data availability

PARIS2 data used for RNA duplex validation were obtained from the NCBI Gene Expression Omnibus (GEO: https://www.ncbi.nlm.nih.gov/geo/) under accession GSE149493. Corresponding processed duplex data for the HEK293T enriched-mRNA dataset HS-HEK293T_Amo-0.5_T4-24h_exo-0h_mRNA were obtained from the CRIS database. icSHAPE data for H9, HeLa, HepG2 and K562 cells were obtained from GEO under accession GSE145805; HEK293T icSHAPE-MaP data were obtained under accession GSE146952.

The RNA-seq datasets used to investigate cell-cycle progression and responses to drug treatment are publicly available through GEO. Corresponding gene- and transcript-level expression data were retrieved through the GEO RNA-seq Experiments Interactive Navigator (GREIN) platform (https://www.ilincs.org/apps/grein/). Gene expression signatures used in the iLINCS analyses were downloaded from the iLINCS database (https://www.ilincs.org). The *Escherichia coli* datasets were obtained directly from GEO. Complete GEO accession numbers, and iLINCS signature identifiers are provided in the main text and in Supplementary Table 3. Processed data supporting the findings of this study are deposited in Zenodo at: https://zenodo.org/records/17363927?preview=1&token=eyJhbGciOiJIUzUxMiJ9.eyJpZCI6IjRiNmY4YTk0LWI5ZWYtNDhjNy05MTRlLTNlYzQ1ZTRiNzE5YiIsImRhdGEiOnt9LCJyYW5kb20iOiJmNjE1YTQxMzkxZWQ3ODVlMDQ1ZDM4ZjhmNGRlZjE0YiJ9.u4SvYzmHKrGQxqc-sREQozT1h1iaX5z7CLcXX1fBRs5dEnsEtLExsxZki201wIDt_WHSqBnsPc8AVlc7KtIkcw

## Code availability

Computational code used in this study is freely available at GitHub https://github.com/Arcadi2020/RNA_thermodynamics_paper, with additional files archived on Zenodo: https://zenodo.org/records/17363927?preview=1&token=eyJhbGciOiJIUzUxMiJ9.eyJpZCI6IjRiNmY4YTk0LWI5ZWYtNDhjNy05MTRlLTNlYzQ1ZTRiNzE5YiIsImRhdGEiOnt9LCJyYW5kb20iOiJmNjE1YTQxMzkxZWQ3ODVlMDQ1ZDM4ZjhmNGRlZjE0YiJ9.u4SvYzmHKrGQxqc-sREQozT1h1iaX5z7CLcXX1fBRs5dEnsEtLExsxZki201wIDt_WHSqBnsPc8AVlc7KtIkcw

