## Supplementary Information for "Evidence of a conserved thermodynamic signature of growth state in living systems"

**Coarse-grained stochastic thermodynamic model of cellular configurational entropy and transcriptome-level structural-stability reweighting**

This model formalizes the physical rationale introduced in the main text. It specifies the assumptions under which cellular-state transitions may alter intracellular configurational entropy and formulates a testable hypothesis linking these changes to transcriptome-level structural-stability reweighting.

**1	Cellular configurational entropy in intracellular nonequilibrium dynamics**

The configurational entropy of the intracellular ensemble is defined as

$$S_{\mathrm{conf}}(t)=-k_{B}\sum_{\Gamma} P(\Gamma,t) \ln P(\Gamma,t),$$

(1)

which quantifies the statistical spread of the probability distribution $P(\Gamma,t)$ over the accessible configurational states $\Gamma$ and $k_{B}$ is the Boltzmann constant.

Time-dependent changes in the intracellular probability distribution $P(\Gamma,t)$ produce corresponding changes in $S_{\mathrm{conf}}$. In an open cellular system, the time evolution of $S_{\mathrm{conf}}$ depends on the balance between irreversible entropy production associated with intracellular stochastic dynamics and net outward entropy flow across the cellular boundary. The latter is not part of $S_{\mathrm{conf}}$ itself but enters its balance through entropy exchange with external reservoirs.

This decomposition provides a thermodynamic framework for interpreting how intracellular organization evolves during processes such as cell-cycle progression or pharmacological perturbation.

### **1.1 Entropy balance**

The evolution of the configurational entropy follows directly from the master equation. Differentiating equation (1) with respect to time gives

$$\frac{dS_{\mathrm{conf}}}{dt}=-k_{B}\sum_{\Gamma} \frac{\partial P(\Gamma,t)}{\partial t}\ln P(\Gamma,t),$$

(2)

where the normalization condition $\sum_{\Gamma} \frac{\partial P(\Gamma,t)}{\partial t}=0$ has been used to eliminate the additional term arising from differentiating $P(\Gamma,t)lnP(\Gamma,t)$.

The time evolution of the probability distribution is governed by the master equation

$$\frac{\partial P(\Gamma,t)}{\partial t}=\sum_{\Gamma^{'}} \left[ W(\Gamma|\Gamma^{'}) P(\Gamma^{'},t)-W(\Gamma^{'}|\Gamma) P(\Gamma,t) \right],$$

(3)

where $W(\Gamma^{'}|\Gamma)$ denotes the transition rate from configuration $\Gamma$ to configuration $\Gamma^{'}$.

Substituting equation (3) into equation (2) yields the standard entropy balance relation of stochastic thermodynamics:

$$\frac{dS_{\mathrm{conf}}}{dt}=\Pi-\Phi,$$

(4)

where $\Pi$ is the irreversible entropy production generated by intracellular stochastic processes and $\Phi$ denotes the entropy flow exchanged with the environment.

Equation (4) represents the nonequilibrium analogue of the second law for stochastic systems: changes in configurational entropy arise from the interplay between internal irreversibility and entropy transfer across the system boundary.

### **1.2 Irreversible entropy production**

The rate of entropy production is

$$\Pi=\frac{k_{B}}{2}\sum_{\Gamma,\Gamma^{'}} J(\Gamma,\Gamma^{'})ln\left[ \frac{W(\Gamma^{'}|\Gamma) P(\Gamma,t)}{W(\Gamma|\Gamma^{'}) P(\Gamma^{'},t)} \right],$$

(5)

where the net probability current between configurations is

$$J(\Gamma,\Gamma^{'})=W(\Gamma^{'}|\Gamma) P(\Gamma,t)-W(\Gamma|\Gamma^{'}) P(\Gamma^{'},t).$$

(6)

The quantity $\Pi$ is the entropy-production rate associated with intracellular stochastic dynamics.

Its non-negativity,

$$\Pi\geq0,$$

is a direct consequence of the second law of thermodynamics.

### **1.3 Entropy exchange with the environment**

The entropy flow term $\Phi$ accounts for entropy transfer between the intracellular system and its surroundings. With the outward-positive convention used here, $\Phi\left( t \right)>0$ denotes net entropy exported from the cell. Under the local-detailed-balance assumption, the logarithmic ratio of the forward and reverse transition rates represents the entropy transferred to the external reservoirs, in units of $k_{B}$, during the transition $\Gamma\to\Gamma^{'}$. Accordingly, the total outward entropy-flow rate is

$$\Phi\left( t \right)=\frac{k_{B}}{2}\sum_{\Gamma,\Gamma^{'}} J(\Gamma,\Gamma^{'})ln\left[ \frac{W(\Gamma^{'}|\Gamma)}{W(\Gamma|\Gamma^{'})} \right]$$

(6a)

Separating the logarithm in the entropy-production expression in equation (5) into its transition-rate and probability-ratio contributions identifies the former with equation (6a) and the latter with $({dS}_{conf})/dt$, thereby recovering the entropy balance in equation (4).

For interpretive purposes, the environmental entropy flow resolved by the model is decomposed as:

$$\Phi\left( t \right)=\Phi^{heat}\left( t \right)+ \Phi^{chem}\left( t \right).$$

(7)

Here, $\Phi^{heat}$ represents the entropy carried by heat dissipated to the environment. Under the isothermal approximation, where the intracellular temperature is effectively clamped by the surrounding medium, the thermal entropy-flow term reduces to $\Phi^{heat}=\dot{Q}_{out} /T$. This expression follows the standard stochastic-thermodynamic treatment of open systems in contact with a thermal reservoir, in which heat exchange contributes directly to entropy flow without requiring an explicit heat-capacity term.

$\Phi^{chem}$ is the chemical-reservoir component associated with exchange of lactate, ions including H^+^, CO_2_, metabolites and other molecular species; its magnitude depends on the corresponding exchange fluxes and reservoir chemical potentials.

At this coarse-grained level, the entropy-production and entropy-flow terms are modelling quantities and are not estimated directly from the transcriptomic data.

**1.4	Initial proliferative non-equilibrium steady state**

Before pharmacologic treatment, we assume an approximately stationary proliferative non-equilibrium state. Within the idealized model, we take the initial state as stationary:

$$\frac{dS_{\mathrm{conf}}}{dt}\simeq0, \Pi^{(0)}\simeq\Phi^{(0)}>0.$$

(8)

The entropy of the cellular state distribution is approximately constant. Entropy continues to be generated internally and exported continuously.

**1.5	Cellular-state transitions**

Processes such as cell-cycle progression or drug treatment modify the transition rates

$$W(\Gamma^{'}|\Gamma),$$

thereby altering probability currents, entropy production $\Pi$ and entropy export $\Phi$.

In this framework, each biological condition corresponds to a distinct stochastic dynamical regime:

$$P(\Gamma,t)\to P_{\alpha}(\Gamma,t),$$

where the index $\alpha$ labels the physiological state (e.g., proliferative phases, drug-induced arrest or recovery).

These states should not be interpreted as different thermodynamic equilibria. Rather, they represent nonequilibrium steady or transient distributions generated by altered intracellular kinetics.

**1.5.1	Drug-induced perturbation**

For perturbations induced by cytostatic molecularly targeted anticancer agents considered here, the transition-rate matrix changes from the initial proliferative state to the drug-exposed state:

$$W^{\left( 0 \right)}(\Gamma^{'}|\Gamma)\to W^{\left( D \right)}(\Gamma^{'}|\Gamma,t),$$

We define changes relative to the initial proliferative state as:

$$\delta\Pi\left( t \right)=\Pi\left( t \right)-\Pi^{\left( 0 \right)}, \delta\Phi\left( t \right)=\Phi\left( t \right)-\Phi^{(0)},$$

(9)

Using the stationary initial balance in equation (8) and the definitions in equation (9), the entropy balance in equation (4) becomes

$$\frac{dS_{\mathrm{conf}} (t)}{dt}=\delta\Pi\left( t \right)-\delta\Phi\left( t \right).$$

(10)

Using the decomposition in equation (7), the same balance can be written explicitly as

$$\frac{dS_{\mathrm{conf}} (t)}{dt}=\delta\Pi\left( t \right)-{\delta\Phi}^{heat}\left( t \right)- {\delta\Phi}^{chem}\left( t \right),$$

(10a)

where ${\delta\Phi}^{heat}$ and ${\delta\Phi}^{chem}$ are defined analogously relative to the initial state.

The general condition for a transient increase in intracellular configurational entropy is therefore

$$\delta\Pi\left( t \right)-\delta\Phi\left( t \right)>0.$$

(11)

In the biological setting considered in this study, growth arrest can reduce internal irreversible entropy production as well as the thermal and chemical-reservoir components of entropy export. The relevant question is therefore not the behavior of any single metabolic readout, but whether the total entropy exported from the cell decreases more or less than the internal entropy production. For the early growth-arrest transitions considered here, we assume that both $\delta\Pi\left( t \right)$ and $\delta\Phi\left( t \right)$ are negative relative to the proliferative baseline:

$$\delta\Pi\left( t \right)<0, \delta\Phi\left( t \right)<0.$$

A transient increase in configurational entropy can arise when the decrease in total environmental entropy flow is greater than the decrease in irreversible entropy production.

$$\left| \delta\Phi\left( t \right) \right|>\left| \delta\Pi\left( t \right) \right|.$$

(12)

A biologically plausible route to this condition is a pronounced early reduction in the chemical-reservoir component associated with growth-linked lactate/H^+^ and metabolite exchange, while contemporaneous changes in thermal entropy flow and irreversible entropy production are comparatively smaller, as explained in Section 1.5.2. Combining equations (10) and (12) gives the central prediction

$$\frac{dS_{\mathrm{conf}}\left( t \right)}{dt}>0.$$

(13)

Thus, during the drug-induced transition from the initial proliferative state toward proliferative arrest, the probability distribution over intracellular configurations broadens while the inequality in equation (12) holds.

**1.5.2	Metabolic evidence motivating the entropy-flow inequality in response to molecularly targeted anticancer agents**

Cellular energy metabolism is supported principally by glycolysis and mitochondrial oxidative phosphorylation, with glycolysis additionally supplying carbon intermediates for biosynthetic pathways. Extracellular acidification rate (ECAR) and oxygen consumption rate (OCR) are widely used experimental readouts for profiling these programmes. ECAR qualitatively reflects glycolysis-associated lactate/H⁺ export, whereas OCR reports oxygen consumption associated predominantly with mitochondrial respiration^1^.

Within the model, total outward entropy flow is decomposed into thermal and chemical-reservoir contributions,$\Phi^{heat}+ \Phi^{chem}$ where the former arises from heat dissipation and the latter from exchange of metabolites, ions, and other molecular species with external reservoirs. The thermal component $\Phi^{heat}$accounts for entropy transfer associated with heat dissipation. In an isothermal cellular environment, heat flow contributes to entropy export solely through $\dot{\frac{Q}{T}}$ , consistent with the standard reservoir-coupling formulation of stochastic thermodynamics.

Instead, the chemical-reservoir component $\Phi^{chem}$ represents entropy carried through the exchange of metabolites, gases, ions and other molecular species.

Among these exchanges, glycolysis-associated lactate/H⁺ export contributes to $\Phi^{chem}$ and is qualitatively reflected by ECAR. A decrease in ECAR, when accompanied by reduced lactate output, is consistent with reduced glycolysis-associated exchange.

In response to cytostatic molecularly targeted agents, growth-associated glycolytic and anabolic programs in human cancer cells are suppressed, whereas mitochondrial oxidative activity is comparatively preserved or adaptively maintained.

This metabolic asymmetry has been consistently observed across multiple cell lines and targeted agents:

1. In FGFR2-fusion-positive intrahepatic cholangiocarcinoma models, the FGFR inhibitor infigratinib reduced glucose uptake, extracellular lactate and ECAR while preserving basal and maximal OCR^2^.
2. In BRAF^V600^-mutant melanoma cells, the BRAF inhibitor vemurafenib induces pronounced suppression of glucose uptake, lactate production and ECAR with comparatively minimal changes in OCR^3^.
3. In human ACHN renal cell carcinoma cells, the JAK1/2 inhibitor momelotinib and the AKT inhibitor MK2206 suppressed glucose uptake, lactate production and ECAR while maintaining mitochondrial OXPHOS through autophagy-supported fatty-acid oxidation^4^.
4. In HER2-positive BT-474 breast cancer cells, the dual HER2/EGFR inhibitor lapatinib induced an immediate decrease in ECAR, whereas OCR remained unchanged over the measured interval^5^.
5. In EGFR-mutant PC-9 lung adenocarcinoma cells, the EGFR inhibitors gefitinib and erlotinib reduced glucose consumption, lactate accumulation and ECAR, whereas OCR remained unchanged^6^.
6. In BCR-ABL1-positive K562, BV173 and KU812 chronic myeloid leukaemia cells, 24 h treatment with imatinib, dasatinib, nilotinib or asciminib markedly attenuated ECAR while producing only a slight reduction in OCR. Glycolytic ATP production and extracellular lactate also declined more strongly than mitochondrial ATP production^7^.
7. In RB-proficient HepG2 hepatocellular carcinoma cells, palbociclib reduced ECAR after 24 h without altering basal or ATP-linked respiration, maximal respiration or spare respiratory capacity^8^.
8. Across different patient-derived glioblastoma models, panobinostat, vorinostat and romidepsin suppressed glycolytic programs. Direct extracellular-flux analyses with panobinostat and vorinostat showed reduced ECAR and increased OCR, while complementary analyses linked the compensatory OXPHOS response to fatty-acid oxidation^9^.
9. In cetuximab-sensitive HN5 and FaDu head-and-neck squamous-cell carcinoma cells, cetuximab reduced ECAR and increased OCR^10^.

Collectively, these studies support a recurrent early-response metabolic asymmetry characterised by stronger or earlier suppression of glycolysis-associated acidification than mitochondrial respiration, which serves in the model as the principal example of a metabolic process that continues to contribute to internal irreversible entropy production ($\Pi$) during early growth arrest. The magnitude of this asymmetry varies with cellular context, tumour type and time after treatment. Comparatively preserved mitochondrial respiration is consistent with the continued energetic requirements of viable growth-arrested cells, including proteostasis, molecular repair, transmembrane ion-gradient regulation and stress response programs^11^.

Although ECAR and OCR do not directly quantify total outward entropy flow ($\Phi$) and internal irreversible entropy production ($\Pi$), respectively, their experimentally observed differential response provides the physiological basis for the coarse-grained thermodynamic representation introduced in Section 1.5.1 and motivation for adopting the sufficient condition in equation (12).

Bacterial adaptation to nutrient limitation provides a distinct metabolic context in which to consider how the same entropy-flow inequality could be satisfied, as discussed in the following section.

**1.5.3	Metabolic evidence motivating the entropy-flow inequality during bacterial nutrient limitation**

An analogous, although metabolically distinct, separation can arise during bacterial adaptation to nutrient limitation. Biomass formation and growth-associated exchange fluxes contract, whereas non-growth-associated carbon catabolism and energy-generating activity can persist. The feature shared with growth arrested cancer cells is therefore the differential attenuation of growth-associated chemical exchange and maintenance-associated energy metabolism.

As no single bacterial measurement is directly equivalent to the ECAR–OCR comparison used in mammalian cells, metabolic changes are assessed by integrating measurements of substrate uptake, biomass formation, metabolite excretion, oxygen consumption and adenylate energy status^12^.

1. Following an abrupt glucose-to-fumarate transition, most *E. coli* cells entered a non- or slow-growing persister state. Absolute fumarate uptake, oxygen consumption and CO₂ production were lower than in fumarate-growing cells; however, per mole of fumarate consumed, persisters produced 5.2-fold less biomass and consumed 2.1-fold more oxygen. Flux-balance analysis further estimated an 8.4-fold greater maximal ATP yield beyond biomass requirements, indicating preferential allocation of the limited substrate toward respiratory energy generation and cellular maintenance rather than growth^13^.
2. Following complete exhaustion of ammonium or sulfate in glucose-containing medium, *E. coli* BW25113 entered stationary phase but continued to consume glucose for more than 24 h. This sustained substrate uptake provides quantitative evidence that nitrogen- or sulfur-limited growth arrest remains coupled to non-growth-associated carbon catabolism, consistent with continued energy production for cellular maintenance rather than biomass formation^14^.

Collectively, these studies provide quantitative evidence that nutrient limitation can separate biomass formation from non-growth-associated energy metabolism: growth-linked fluxes contract, whereas carbon catabolism, respiratory activity and ATP-generating capacity can persist. This is conceptually analogous to the preferential suppression of glycolysis-associated acidification relative to mitochondrial respiration in human cancer cells in response to drug-induced growth arrest (Section 1.5.2).

Within the model, contraction of growth-linked substrate and metabolite exchange is expected to reduce the chemical-reservoir component of outward entropy flow, whereas retained maintenance-associated energy metabolism indicates that internal entropy production does not decline proportionately. When the reduction in total outward entropy flow exceeds that in internal entropy production, the sufficient condition in equation (12) is satisfied and intracellular configurational entropy is predicted to increase transiently. These metabolic measurements do not quantify either thermodynamic term directly; rather, they establish the biological plausibility of the proposed entropy-flow imbalance during bacterial adaptation to nutrient limitation.

**1.5.4	Integrated transient response**

Over an adaptive interval $0<t<\tau$, the cumulative change in intracellular configurational entropy is

$$\Delta S_{\mathrm{conf}}\left( \tau\right)=S_{\mathrm{conf}}\left( \tau\right)- S_{\mathrm{conf}}\left( 0 \right)=\int_{0}^{\tau} \left[ \delta\Pi\left( t \right)-\delta\Phi\left( t \right) \right]dt.$$

(14)

A net increase in configurational entropy over this interval occurs when

$$\int_{0}^{\tau} \left[ \delta\Pi\left( t \right)-\delta\Phi\left( t \right) \right]dt>0.$$

(15)

The integrated condition is more general than requiring the instantaneous condition in equation (11) to hold at every time point. Thus, brief intervals in which entropy export does not decline more strongly than irreversible entropy production do not preclude a net transient increase in configurational entropy over the adaptive response.

As arrested cells approach a new nonequilibrium steady state, ${dS}_{\mathrm{conf}}/dt$ tends to zero, by equation (4),

$$\frac{dS_{conf}(t)}{dt}\to0,$$

(16)

By equation (4), the new nonequilibrium steady state satisfies

$$\Pi^{(arrest)}\simeq\Phi^{(arrest)}$$

(17)

Thus, the model predicts transient increase in intracellular configurational entropy, not unlimited accumulation of configurational entropy.

**1.5.5	Cell-cycle-dependent modulation of the entropy balance**

Normal cell-cycle progression can be treated as a phase-dependent modulation of the same stochastic-thermodynamic balance. Unlike cytostatic drug treatment, cell-cycle progression does not impose a sustained transition from proliferation to arrest. Instead, it redistributes metabolic, biosynthetic, replication-associated, and mitotic activities across successive proliferative phases^15-17^. We therefore allow the transition-rate matrix to depend on cell-cycle phase,

$$W(\Gamma^{'}|\Gamma,t)=W^{\phi}(\Gamma^{'}|\Gamma,t), \phi\in\left\{ G_{1} , S , G_{2} , M \right\},$$

(18)

The configurational-entropy balance remains

$$\frac{dS_{conf}^{\left( \phi\right)}(t)}{dt}=\Pi^{\left( \phi\right)}(t)-\Phi^{\left( \phi\right)}(t)$$

(19)

Thus, transitions between cell-cycle phases may transiently broaden or narrow the intracellular configurational distribution depending on the relative modulation of internal entropy production and entropy export. At a given cell-cycle phase, the intracellular configurational distribution broadens when internal irreversible entropy production exceeds entropy export,

$$\Pi^{\left( \phi\right)}(t)-\Phi^{\left( \phi\right)}(t)>0$$

and narrows when entropy export exceeds internal irreversible entropy production,

$$\Pi^{\left( \phi\right)}(t)-\Phi^{\left( \phi\right)}(t)<0$$

This cell-cycle formulation permits phase-specific transient increases and decreases in configurational entropy during continued proliferation, providing a basis for examining whether RNA structural-stability profiles vary systematically across cell-cycle transitions.

Cell-cycle regulators coordinate glycolysis, oxidative phosphorylation and biosynthetic metabolism, with effects varying across cell types and phases^15,18^. In synchronized HeLa cells, isotope tracing and flux modelling revealed complementary changes in glucose- and glutamine-derived TCA-cycle fluxes, with glucose oxidation peaking in late G1 and glutamine metabolism increasing during S phase, while oxygen consumption remained approximately constant^19^. Interestingly, isothermal calorimetry in zebrafish embryos demonstrated heat-flow oscillations dependent on cell-cycle signaling^20^.

These observations motivate phase-dependent metabolic and dissipative fluxes in the model, but do not directly quantify internal entropy production $\Pi$or total entropy export $\Phi$.

**1.6	Transcriptome-level structural-stability projection and abundance reweighting**

Within the model, phase-specific variation in the balance between internal entropy production and entropy export can increase or decrease intracellular configurational entropy during cell-cycle progression (equation (19)), whereas growth arrest can produce a transient increase when the reduction in entropy export exceeds that in internal entropy production (equations (10–15)).

This framework provides the rationale for examining whether such configurational remodelling is reflected in molecular subsystems of the cell. Among these, the transcriptome is particularly informative, as gene expression responds rapidly to physiological and environmental cues.

We therefore asked whether transitions between cellular states differing in proliferative and metabolic activity are accompanied by systematic shifts in abundance-weighted RNA folding stability. This abundance-weighted profile is not treated as a direct measure of cellular configurational entropy, entropy production or entropy export. Instead, it serves as a transcriptome-level structural-stability projection of the predicted intracellular configurational remodelling.

To define this projection, let $i$ index the annotated transcripts, let $j$ be the summation index over all annotated transcripts and let $c$ denote a biological condition. Let $N_{i}^{(c)}$ denote the expression-derived abundance of transcript $i$ in condition $c$. Its normalized abundance weight is defined as

$$w_{i}^{(c)}=\frac{N_{i}^{(c)}}{\sum_{j} N_{j}^{(c)}}$$

Here, $j$ indexes all annotated transcripts, such that

$$\sum_{i} w_{i}^{(c)}=1$$

For each transcript $i$, ${\Delta G}_{fold,i}$ denotes its sequence-derived, length-normalized minimum-free-energy descriptor calculated under fixed folding conditions. This value is invariant across biological conditions; only $N_{i}^{(c)}$ and hence $w_{i}^{(c)}$ changes with condition.

The abundance-weighted mean of predicted ${\Delta G}_{fold}$ in condition $c$ is therefore

$$\overline{\Delta G}_{fold}^{\left( c \right)}=\sum_{i} w_{i}^{(c)}{\Delta G}_{fold,i}$$

For comparisons between conditions, let $r$ denote a reference baseline or untreated condition. The corresponding shift in abundance-weighted predicted folding free energy is

$$\Delta\overline{\Delta G}_{fold}^{\left( c,r \right)}=\overline{\Delta G}_{fold}^{\left( c \right)}-\overline{\Delta G}_{fold}^{\left( r \right)}=\sum_{i} \left[ w_{i}^{\left( c \right)}-w_{i}^{\left( r \right)} \right]\Delta G_{fold,i}$$

(20)

Equation (20) follows by substituting the definitions of the two abundance-weighted means and combining the sums. Each transcript’s $\Delta G_{fold,i}$ value is fixed across conditions; therefore, between-condition differences arise solely from changes in abundance weights. A positive $\Delta\overline{\Delta G}_{fold}^{\left( c,r \right)}$ denotes a shift toward less-negative values, whereas a negative value denotes a shift toward more-negative values.

Across cell-cycle transitions, equation (19) permits phase-specific increases or decreases in intracellular configurational entropy. Accordingly, no universal direction is imposed on the corresponding RNA structural-stability projection during cell-cycle progression. For growth-arrest comparisons, however, $r$ denotes the matched proliferative or actively growing reference condition and $c$ denotes the arrested or growth-restricted condition sampled after the interval $\tau$. The experimental evidence reviewed in Sections 1.5.2 and 1.5.3 motivates the metabolic asymmetry represented by equation (12). Combining equations (10) and (12) gives $dS_{\mathrm{conf}}\left( t \right)/dt>0$ while this inequality holds, and when the cumulative imbalance over the experimental interval is positive, equations (14–15) give

$$\Delta S_{\mathrm{conf}}\left( \tau\right)>0.$$

We therefore formulate the transcriptome-level directional projection hypothesis as

$$\Delta S_{\mathrm{conf}}\left( \tau\right)>0 \Longrightarrow\Delta\overline{\Delta G}_{fold}^{\left( c,r \right)}>0.$$

(21)

A positive $\Delta\overline{\Delta G}_{fold}^{\left( c,r \right)}$corresponds to a shift of the expressed RNA pool toward less-negative predicted folding-free-energy values. Equation (21) specifies the expected direction of the transcriptome-level projection; it is not a thermodynamic identity or a quantitative proportionality between the two quantities. Accordingly, the abundance-weighted RNA descriptor is not interpreted as a direct measure of cellular configurational entropy, entropy production, entropy export or RNA conformational entropy. Rather, equation (21) states the hypothesis that the predicted transient increase in intracellular configurational entropy during growth arrest is accompanied by compositional reweighting of the transcriptome toward RNAs with less-negative fixed $\Delta G_{\mathrm{fold}}$values.

Equation (20) shows that this growth-arrest-associated shift can arise through reduced representation of transcripts with more-negative $\Delta G_{\mathrm{fold}}$values, increased representation of transcripts with less-negative values, or both. The effect is therefore compositional and does not imply that pharmacological treatment or nutrient limitation change the predicted $\Delta G_{\mathrm{fold}}$of any individual transcript. Distinct transcriptional responses can consequently produce the same aggregate direction of reweighting.

**Table 1.** Principal quantities used in the entropy-balance and transcriptome-reweighting framework.

| **Quantity** | **Definition or relation** | **Meaning and scope** |
| --- | --- | --- |
| $S_{\mathrm{conf}}(t)$ | $S_{\mathrm{conf}}(t)=-k_{B}\sum_{\Gamma} P(\Gamma,t) \ln P(\Gamma,t)$ | Gibbs–Shannon configurational entropy of the intracellular state distribution. |
| $\Pi(t)$ | $\Pi(t) \geq0$ | Internal irreversible entropy-production rate of the coarse-grained cellular process. This is a modelling quantity not estimated directly from transcriptomics. |
| $\Phi(t)$ | $\Phi(t) = \Phi^{heat}(t) + \Phi^{chem}(t)$ | Total outward entropy flow, decomposed into thermal and chemical-reservoir components. |
| $\delta\Pi(t); \delta\Phi(t)$ | $\delta\Pi\left( t \right)=\Pi\left( t \right)-\Pi^{\left( 0 \right)}$  $\delta\Phi\left( t \right)=\Phi\left( t \right)-\Phi^{(0)}$ | Changes in entropy production and entropy export relative to the initial proliferative state. |
| $w_{i}^{(c)}$ | $\frac{N_{i}^{(c)}}{\sum_{j}N_{j}^{(c)}}$ | Relative expression-derived abundance weight of transcript i in condition c; the weights sum to one. |
| ${\Delta G}_{fold,i}$ | Sequence-derived, length-normalized minimum-free-energy descriptor calculated under fixed folding conditions. | Fixed, sequence-specific structural-stability descriptor (kcal·mol⁻¹·nt⁻¹); more-negative values indicate greater predicted structural stability. |
| $\overline{\Delta G}_{\mathrm{fold}}^{(c)}$ | $\overline{\Delta G}_{\mathrm{fold}}^{(c)} = \sum_{i}w_{i}^{(c)}{\Delta G}_{fold,i}$ | Normalized abundance-weighted transcriptome-level folding-stability descriptor. |
| $\Delta\overline{\Delta G}_{fold}^{\left( c,r \right)}$ | $\Delta\overline{\Delta G}_{fold}^{\left( c,r \right)}=\overline{\Delta G}_{fold}^{\left( c \right)}-\overline{\Delta G}_{fold}^{\left( r \right)}$  $=\sum_{i} \left[ w_{i}^{\left( c \right)}-w_{i}^{\left( r \right)} \right]\Delta G_{fold,i}$ | Between-condition shift in the abundance-weighted folding-stability descriptor; positive values indicate a less-negative profile and negative values a more-negative profile. |

**Note.** $\Gamma$ indexes accessible intracellular configurations; P($\Gamma,t$) is the probability distribution; $k_{B}$ is the Boltzmann constant; i and j index annotated transcripts; c and r denote the comparison and reference conditions, respectively; $N_{i}^{(c)}$ denotes expression-derived transcript abundance.

**Supplementary Fig. 1**

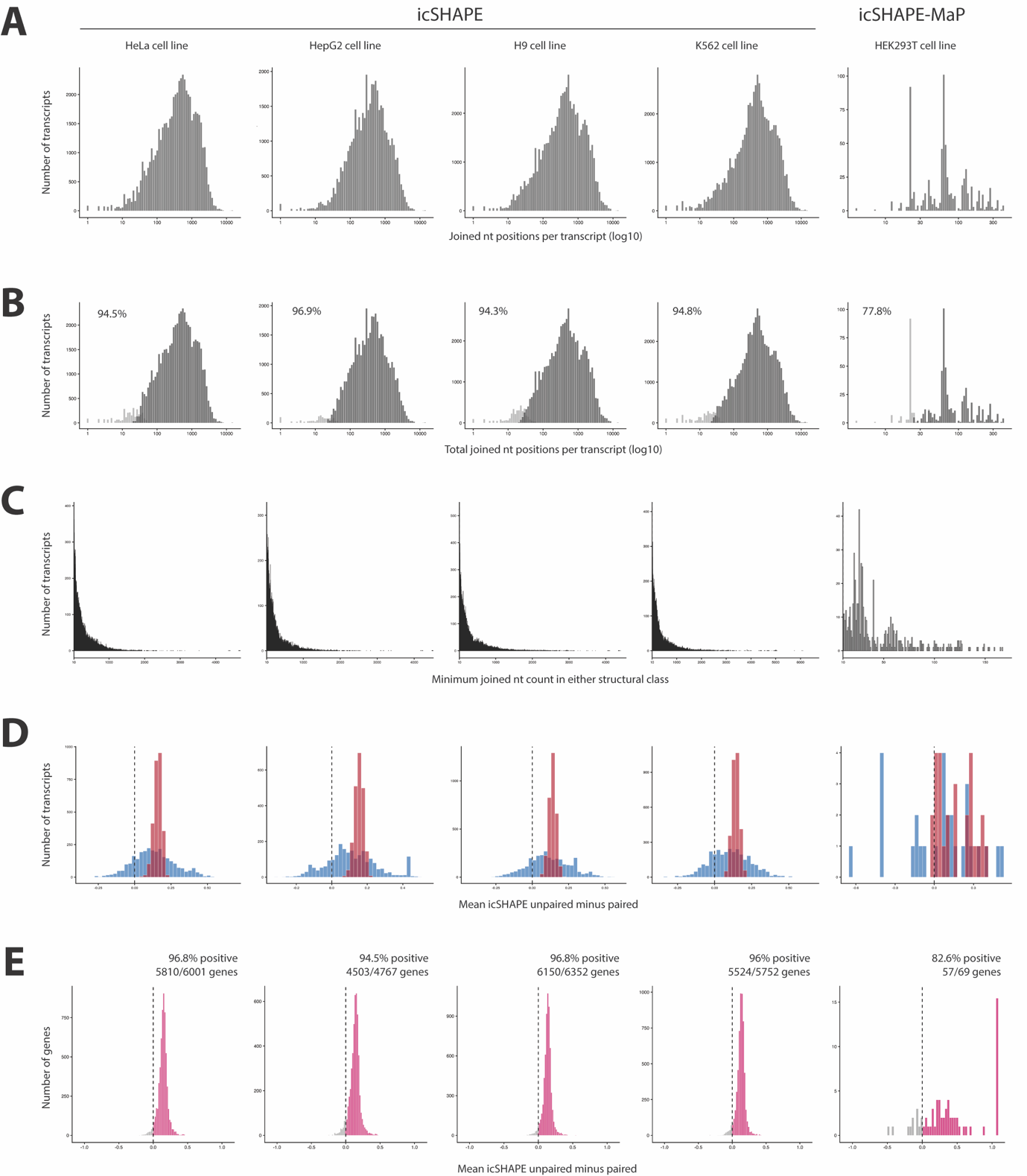

**Supplementary Fig. 1.** **Nucleotide coverage and filtering for the ViennaRNA/icSHAPE concordance analysis.** (**A**) Distribution of joined nucleotide coverage per transcript across HeLa, HepG2, H9 and K562 icSHAPE datasets and the HEK293T icSHAPE-MaP dataset. Joined positions correspond to transcript-mapped nucleotides for which both an experimental icSHAPE/icSHAPE-MaP reactivity score and a ViennaRNA paired/unpaired assignment were available. Distributions are shown before application of the paired/unpaired coverage filter. (**B**) Retention of joined transcripts after paired/unpaired coverage filtering. Transcripts were classified according to whether they passed the coverage criterion used for transcript-level concordance analysis. Light grey bars indicate transcripts that failed the paired/unpaired coverage filter, whereas dark grey bars indicate retained transcripts. Percentages indicate the fraction of joined transcripts retained after filtering. (**C**) Distribution of limiting paired/unpaired coverage among retained transcripts. For each retained transcript, limiting coverage was defined as the lower of the two joined nucleotide counts assigned to the paired and unpaired ViennaRNA classes. (**D**) Coverage-stratified transcript-level concordance analysis. Within each cell line, transcripts were stratified into the lower and upper 5% of the limiting-coverage distribution. For each transcript, concordance was quantified as the mean experimental reactivity at ViennaRNA-predicted unpaired positions minus the mean experimental reactivity at ViennaRNA-predicted paired positions. Positive values therefore indicate higher experimental accessibility at predicted unpaired positions, consistent with concordance between ViennaRNA structural assignments and *in vivo* RNA accessibility. Blue bars indicate the lower 5% coverage group, and red bars indicate the upper 5% coverage group. (**E**) Canonical-transcript collapsed gene-level concordance analysis. For each gene, the Ensembl canonical transcript was retained when available, and concordance was quantified as the mean experimental reactivity at ViennaRNA-predicted unpaired positions minus the mean experimental reactivity at ViennaRNA-predicted paired positions. Positive values indicate genes for which predicted unpaired positions showed higher *in vivo* accessibility than predicted paired positions. Grey bars indicate genes with non-positive differences, whereas pink bars indicate genes with positive differences. Percentages indicate the fraction of genes with positive mean differences in each dataset.

**Supplementary Fig. 2**

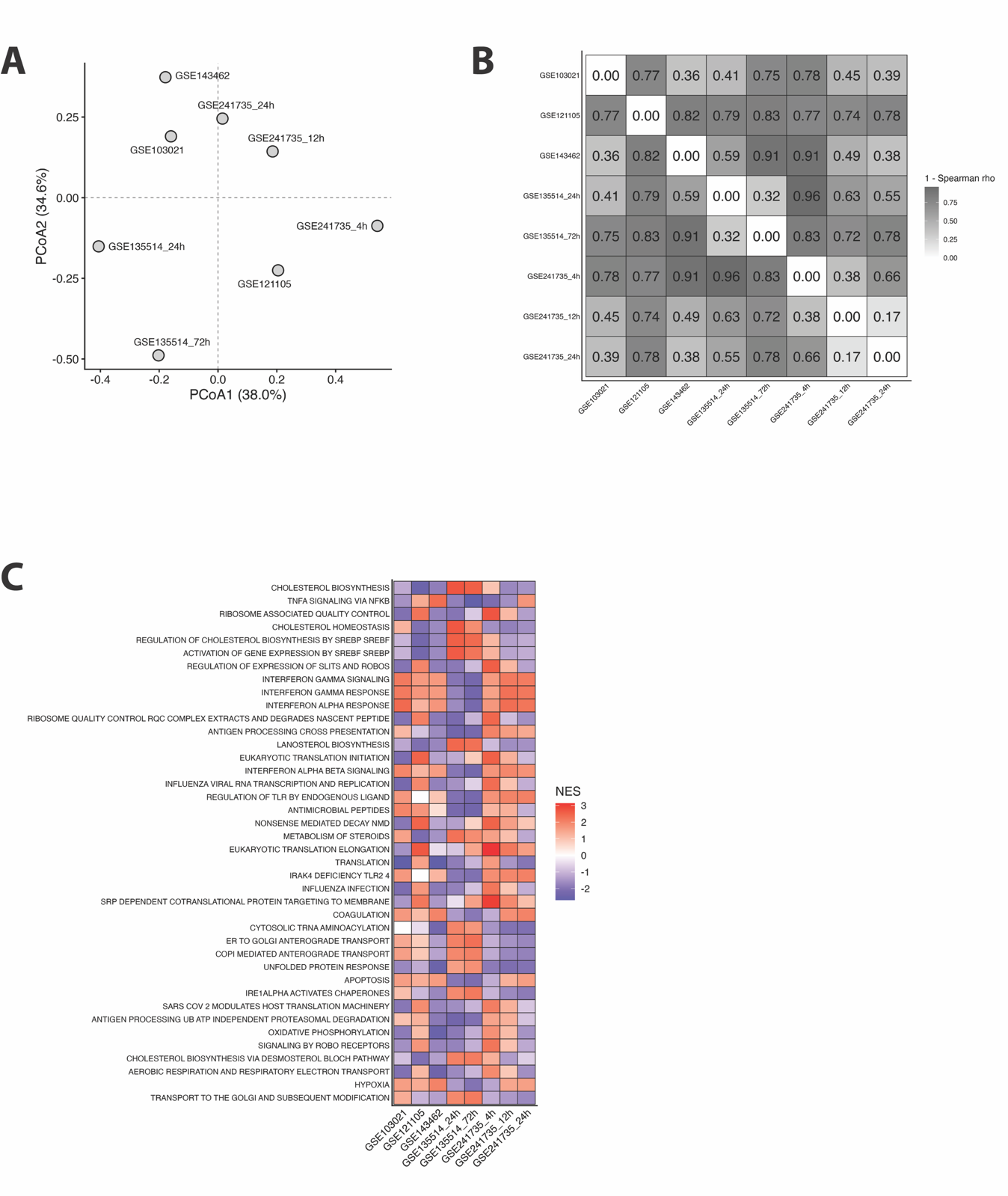

**Supplementary Fig. 2. Distinct pathway-level transcriptional responses accompany a shared RNA structural-stability signature across GREIN drug-response datasets. (A)** Principal-coordinate analysis of GSEA normalized enrichment-score (NES) profiles across GREIN datasets. Each point represents one drug-treated versus untreated control transcriptional contrast. **(B)** Pairwise Spearman-distance heatmap comparing GSEA normalized enrichment-score (NES) profiles across the treatment contrasts. A common pathway set was defined as pathways with a Benjamini–Hochberg-adjusted P value ≤ 0.25 in at least one contrast. Pairwise distances were then calculated as 1 − Spearman’s ρ using the NES values of this common pathway set across all contrasts. Larger values indicate more distinct pathway-level transcriptional profiles. **(C)** Heatmap showing the 40 pathways with the greatest variation in GSEA normalized enrichment scores (NES) across the treatment contrasts. Pathways were selected from the common pathway set defined in panel B and ranked by the standard deviation of their NES values across all contrasts. Blue and red indicate negative and positive enrichment, respectively.

**Supplementary Fig. 3**

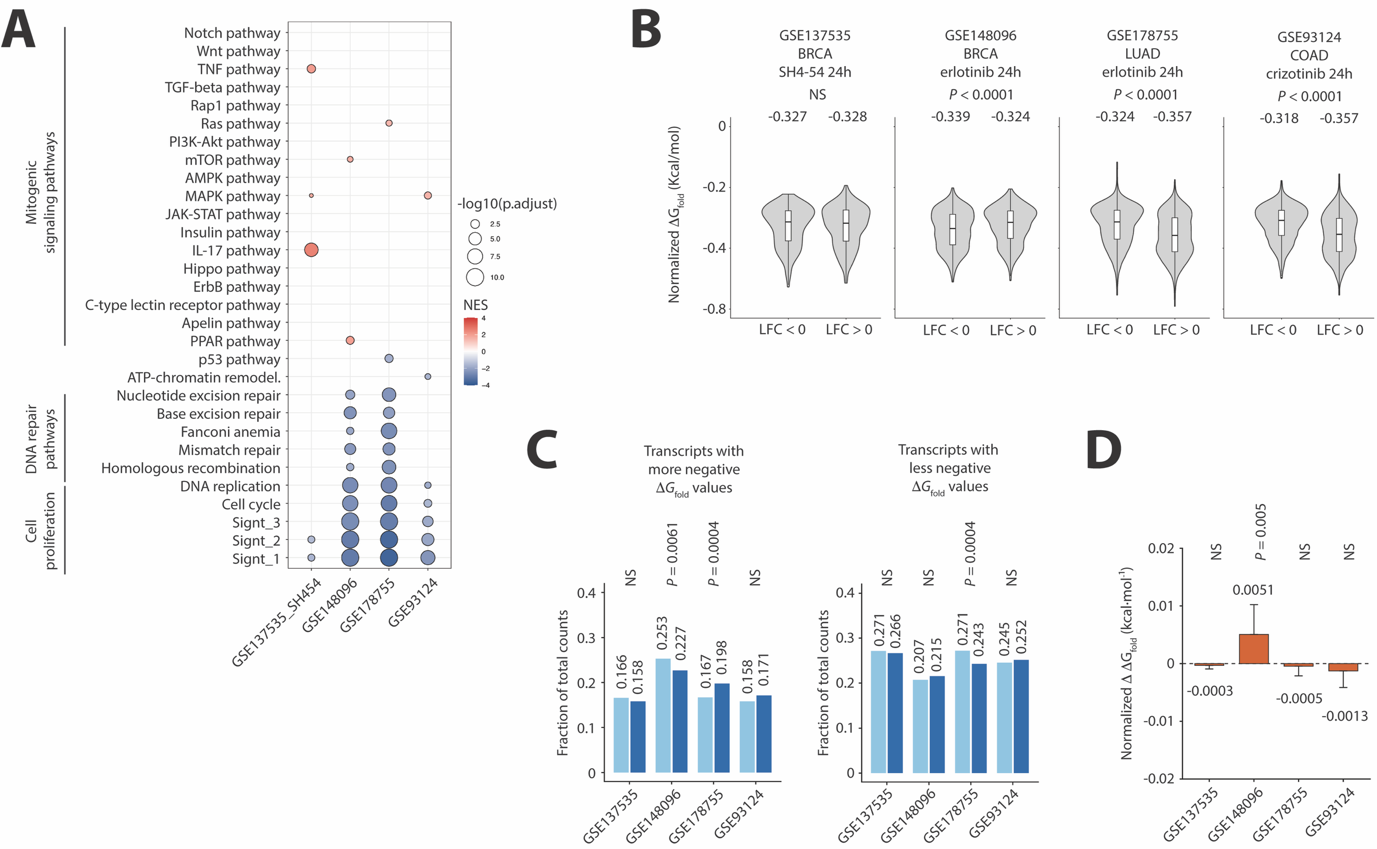

**Supplementary Fig. 3. Residual mitogenic signaling is associated with transcriptome reweighting toward greater predicted RNA structural stability.** (**A**) Gene set enrichment analysis (GSEA) of KEGG pathways in cancer cell lines treated with molecularly targeted agents but exhibiting incomplete suppression of mitogenic signaling. Dot size indicates –log₁₀ adjusted p-value, and color reflects normalized enrichment score (NES). (**B**) Δ*G*_fold_ distributions of significantly up- and downregulated transcripts in the respective cell lines. Unlike conditions in which cells undergo complete arrest of cell proliferation, upregulated transcripts tend to display more negative Δ*G*_fold_ values, indicating increased structural stability. Statistical significance was assessed using the Wilcoxon rank-sum test. (**C**) Fractions of total normalized counts contributed by transcripts in the lower and upper Δ*G*_fold_ quartiles within each significantly differentially expressed transcript pool under control and treatment conditions. *P* values from two-sided folding-label permutation tests (10,000 permutations; BH-adjusted). (**D**) Treatment-minus-control differences in transcriptome-wide TPM-weighted mean normalized Δ*G*_fold_. Error bars show propagated SEM. Negative and positive values indicate shifts toward more- and less-negative Δ*G*_fold_, respectively. *P* values were obtained from 999 permutations of Δ*G*_fold_ across transcripts with TPM contrast weights fixed and were BH-adjusted.

**Supplementary Fig. 4**

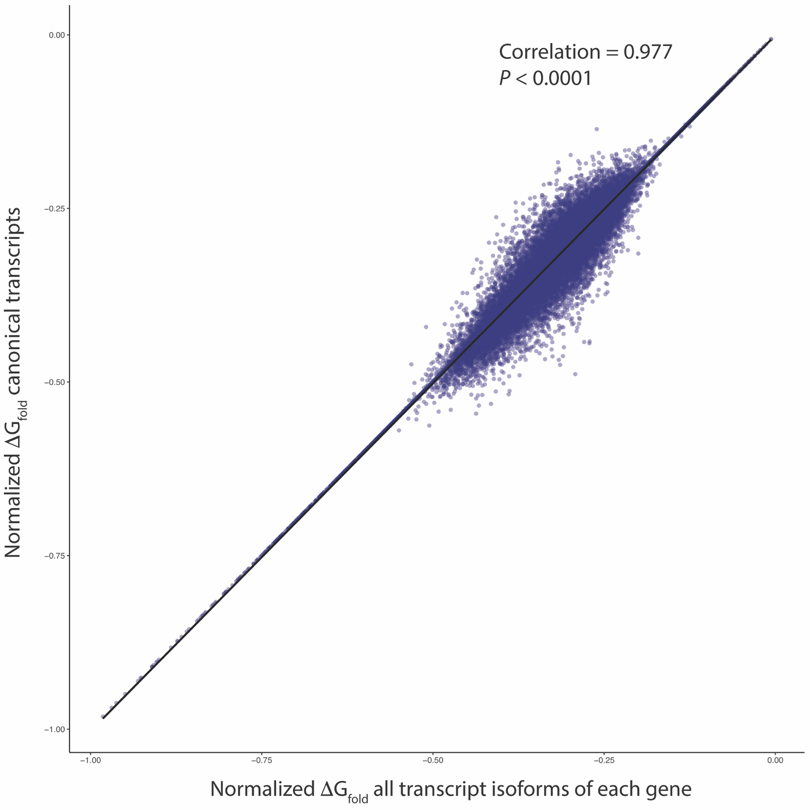

**Supplementary Fig. 4. Canonical transcript Δ*G*_fold_ values provide a reliable approximation of gene-level predicted RNA structural stability.** Scatter plot comparing Δ*G*_fold_ values derived from Ensembl canonical transcripts and the mean Δ*G*_fold_ values computed across all isoforms per gene. The strong correlation (*P* < 0.0001) confirms that canonical transcripts serve as robust proxies for transcriptome-wide thermodynamic analyses in microarray-based datasets.

**Supplementary Fig. 5**

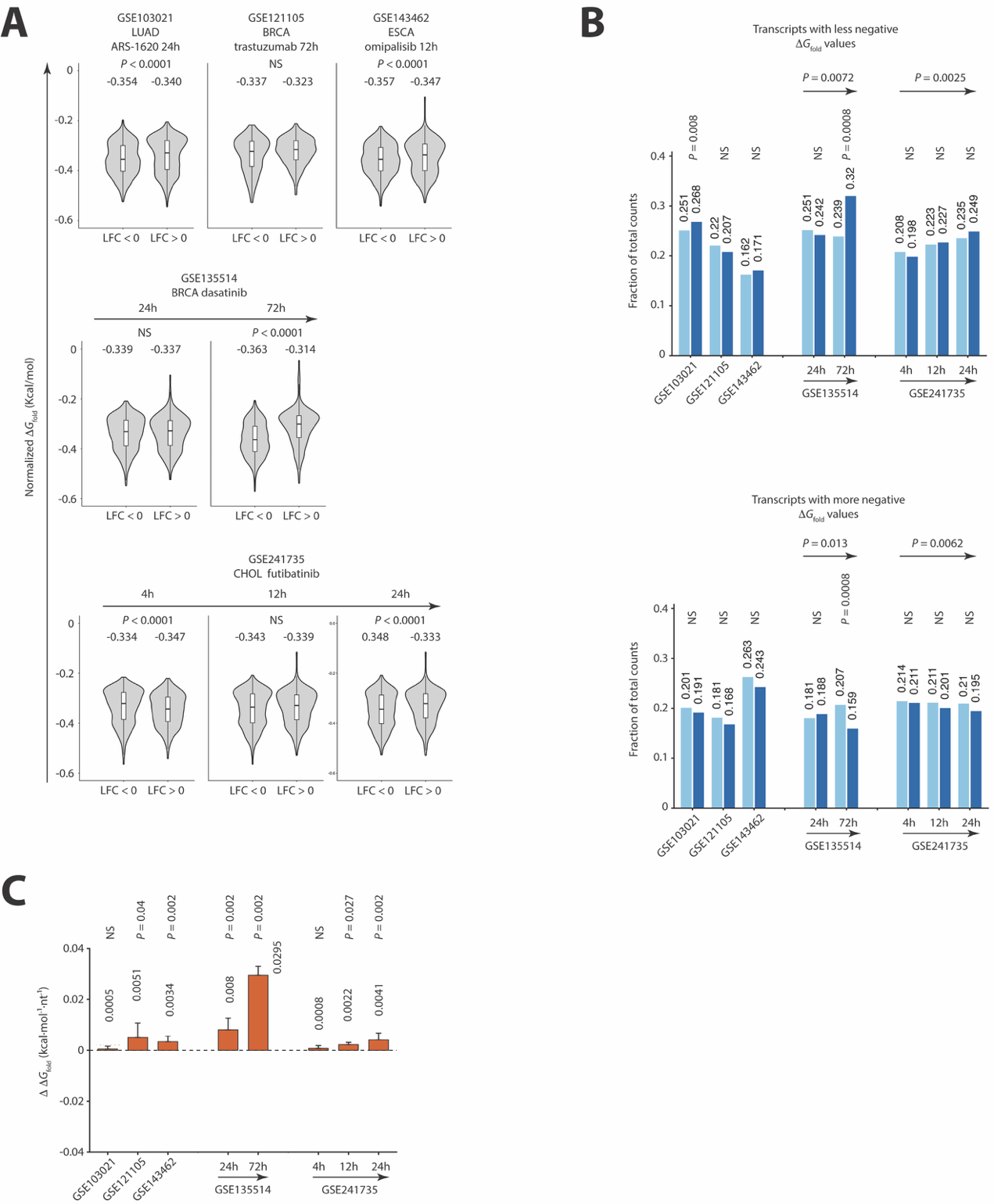

**Supplementary Fig. 5. Canonical-transcript analysis recapitulates thermodynamic RNA structurome profiles obtained using the complete transcript isoform set.** (**A**) Violin plots showing normalized transcript Δ*G*_fold_ values for significantly downregulated and upregulated transcripts following drug treatment. P values from two-sided Wilcoxon rank-sum tests. NS, not significant; ESCC, esophageal squamous-cell carcinoma; BRCA, breast carcinoma; LGC, lung giant-cell carcinoma; CHOL, intrahepatic cholangiocarcinoma; LFC, log2 fold change. **(B)** Fractions of total normalized counts contributed by transcripts in the upper (less-negative Δ*G*_fold_) and lower (more-negative Δ*G*_fold_) quartiles within each significantly differentially expressed canonical transcript pool. Light and dark blue indicate untreated controls and drug-treated samples, respectively. *P* values from folding-label permutation tests (10,000 permutations) for individual comparisons and Welch–Satterthwaite contrasts for temporal changes in treated-minus-control fractions; all two-sided and separately BH-adjusted. **(C)** Transcriptome-wide, transcript abundance-weighted ΔΔ*G*_fold_ index, calculated for the expressed canonical-transcript pool as the drug-treated value minus the untreated-control value. Error bars show propagated SEM; permutation *P* values were Benjamini–Hochberg adjusted across datasets.
**Supplementary Fig. 6**

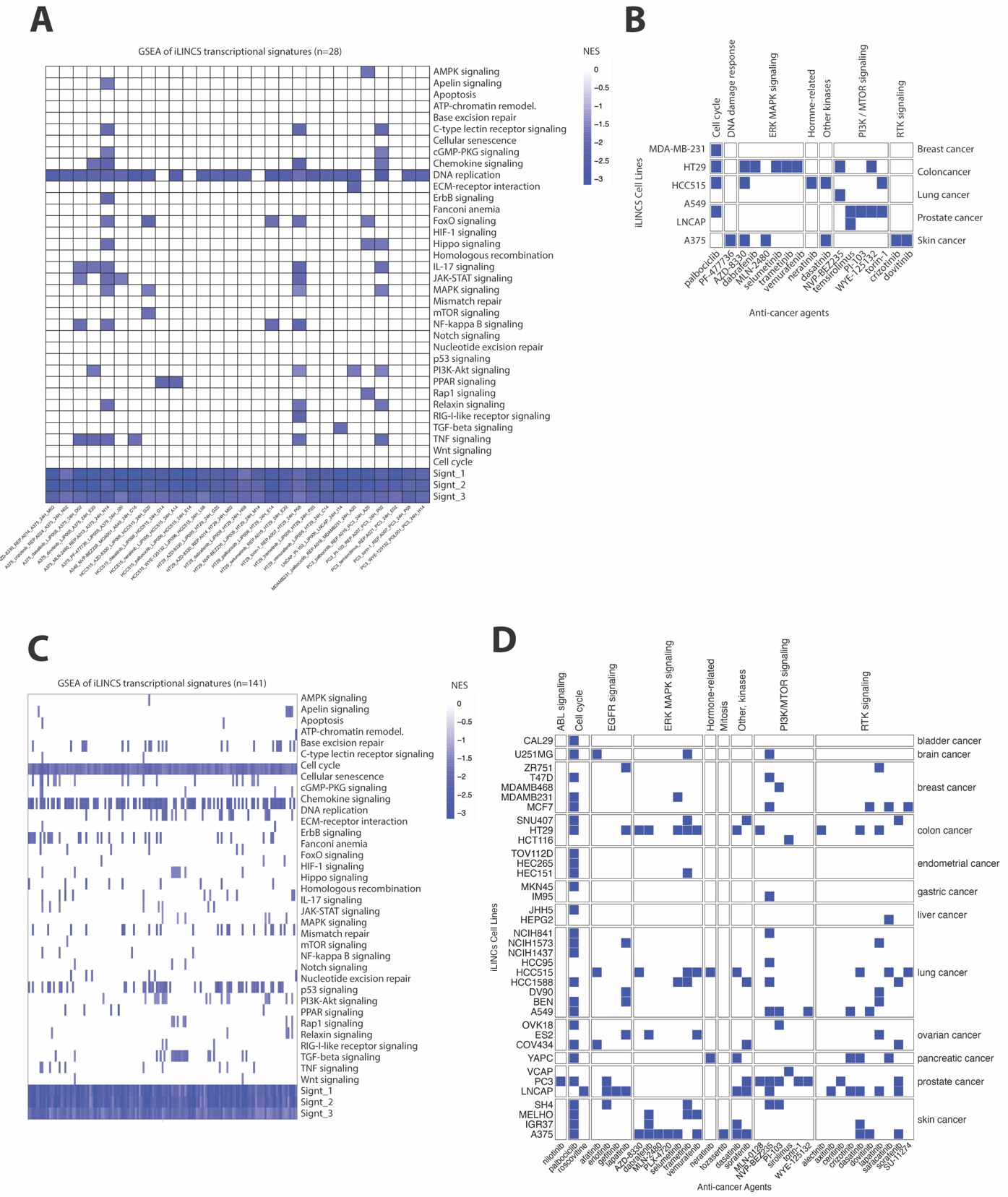

**Supplementary Fig. 6. Identification of cell lines undergoing growth arrest in the iLINCS datasets.** (**A**, **C**) KEGG pathway enrichment analysis demonstrating downregulation of cell cycle and proliferation related pathways in iLINCS cell lines treated with targeted agents at 3.33 µM (**A**) and 10 µM (**C**). (**B**, **D**). Heatmaps depicting selected iLINCS cell lines that meet criteria for complete growth arrest.

**Supplementary Fig. 7**

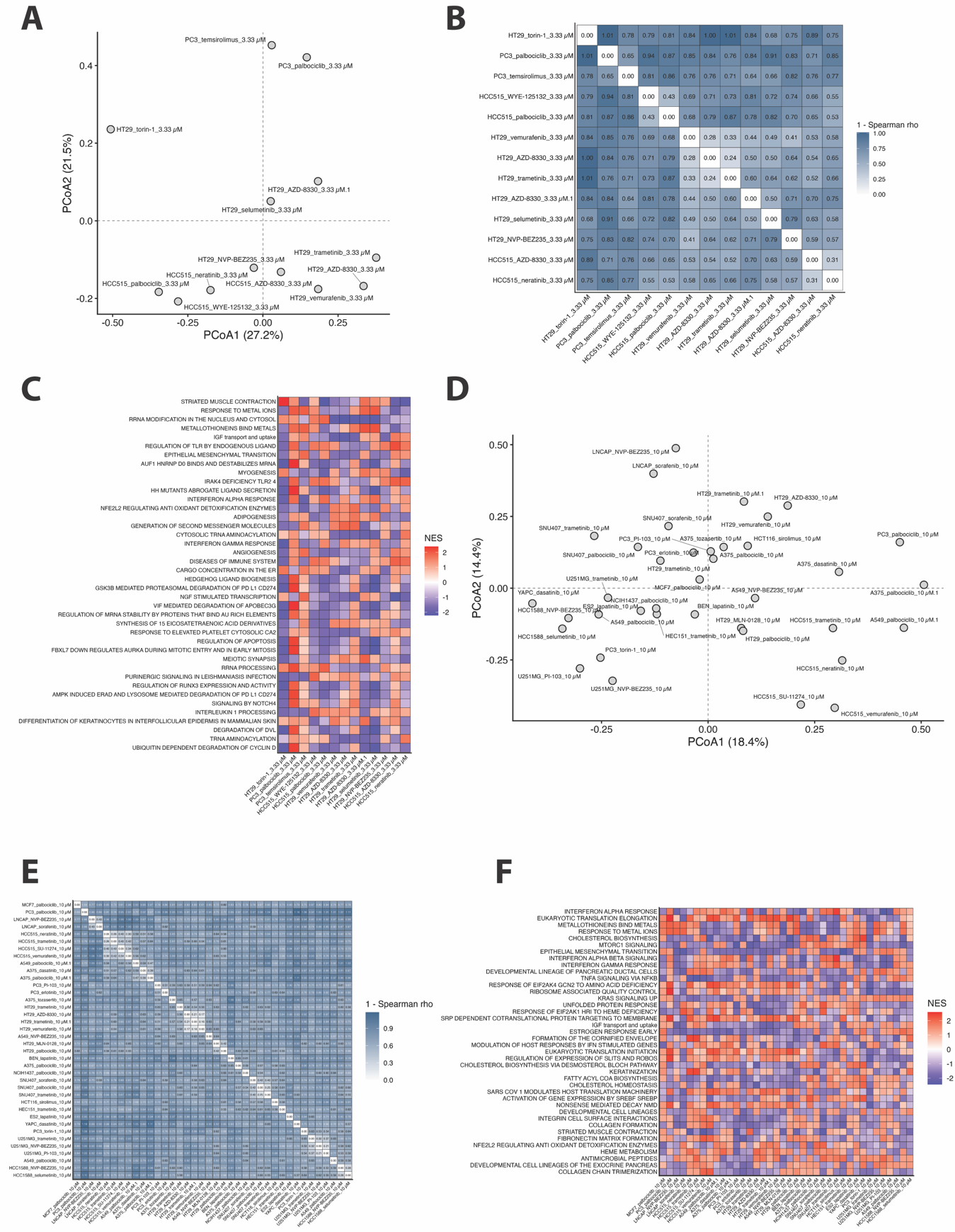

**Supplementary Fig. 7. Distinct pathway-level transcriptional responses accompany a shared RNA structural-stability signature across iLINCS drug-response profiles.** (**A**–**C**) Analysis of iLINCS signatures at 3.33 µM with significant upregulation of transcripts with less negative predicted Δ*G*_fold_ values. (**A**) Principal-coordinate analysis of GSEA normalized enrichment-score (NES) profiles. (**B**) Pairwise Spearman-distance heatmap, calculated as 1 − Spearman’s ρ. (**C**) Heatmap of the pathways with the highest variation in NES values across signatures. (**D**–**F**) Corresponding analysis of iLINCS signatures at 10 µM. (**D**) Principal-coordinate analysis of GSEA/NES profiles. (**E**) Pairwise Spearman-distance heatmap. (**F**) Heatmap of the pathways with the highest variation in NES values across signatures.

**Supplementary Fig. 8**

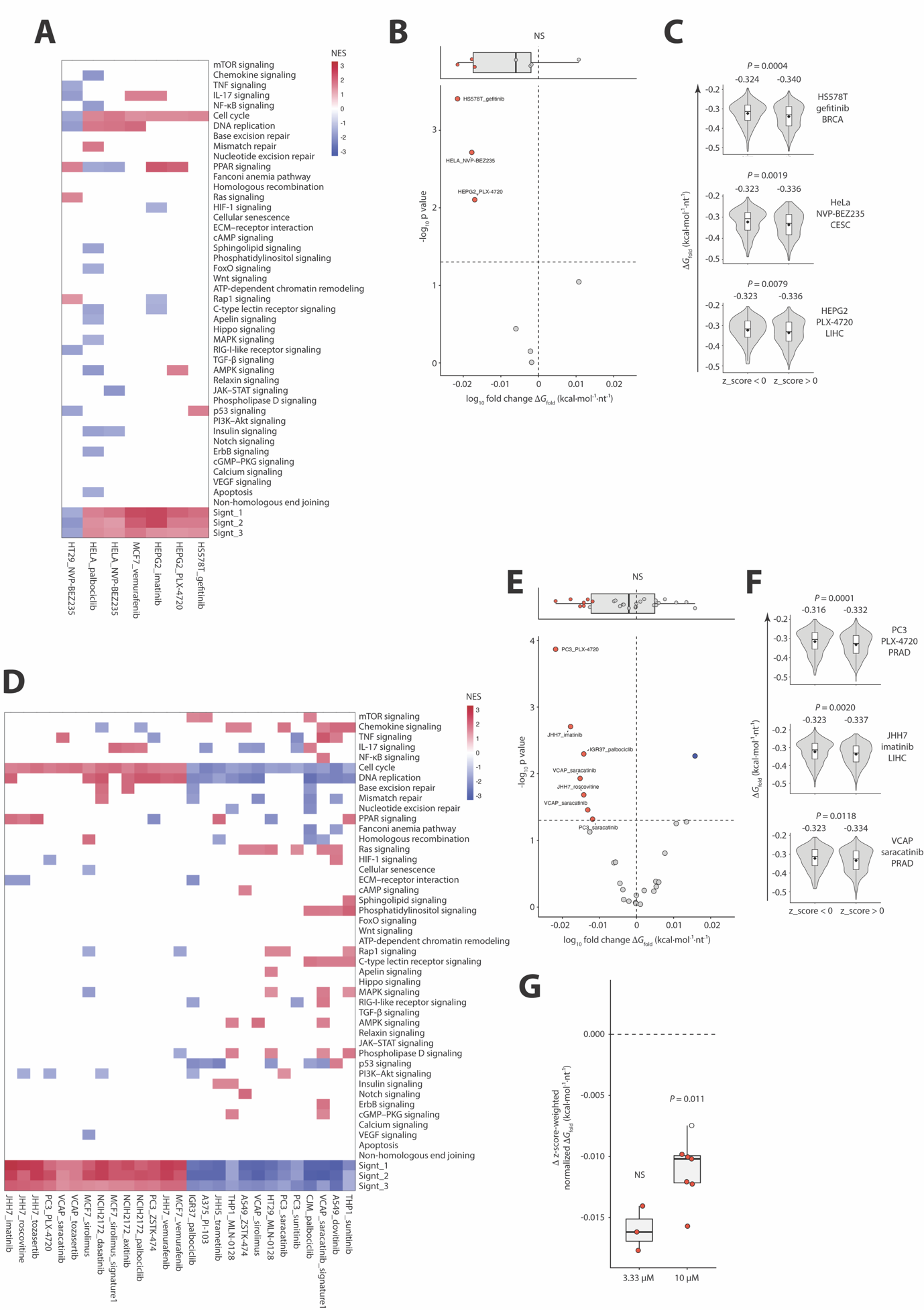

**Supplementary Fig. 8. iLINCS signatures with positive enrichment of proliferative or mitogenic programs are associated with transcriptome reweighting toward greater predicted RNA structural stability.** (**A**)**,** GSEA normalized enrichment score (NES) heatmap for the qualifying 24-h iLINCS signatures at 3.33 µM. (**B**), Distribution and volcano plot of normalized Δ*G*_fold_ shifts across the 3.33 µM signatures. The shift was defined as log₁₀(mean normalized Δ*G*_fold_ of negatively perturbed genes/mean normalized Δ*G*_fold_ of positively perturbed genes); negative values indicate preferential upregulation of transcripts with more negative Δ*G*_fold_. Red, blue and grey denote significant more-negative shifts, significant less-negative shifts and nonsignificant signatures, respectively. Dashed lines indicate zero shift and *P* = 0.05. (**C**), Normalized Δ*G*_fold_ distributions of positively and negatively perturbed genes in three representative 3.33 µM signatures with significant more-negative shifts. (**D**), GSEA NES heatmap for qualifying 24-h iLINCS signatures at 10 µM. **E**, Distribution and volcano plot of normalized Δ*G*_fold_ shifts across the 10 µM signatures, displayed as in B. (**F**), Normalized Δ*G*_fold_ distributions in three representative 10 µM signatures with significant more-negative shifts, displayed as in C. (**G**), Z-score-weighted normalized Δ*G*_fold_ perturbation index for signatures significantly upregulating more-negative Δ*G*_fold_ transcripts. Negative values indicate that positively perturbed genes have more negative Δ*G*_fold_ than negatively perturbed genes. Filled points indicate significant signature-level permutation tests; displayed group-level *P* values are from one-sided Wilcoxon signed-rank tests against zero.

**Supplementary Fig. 9**

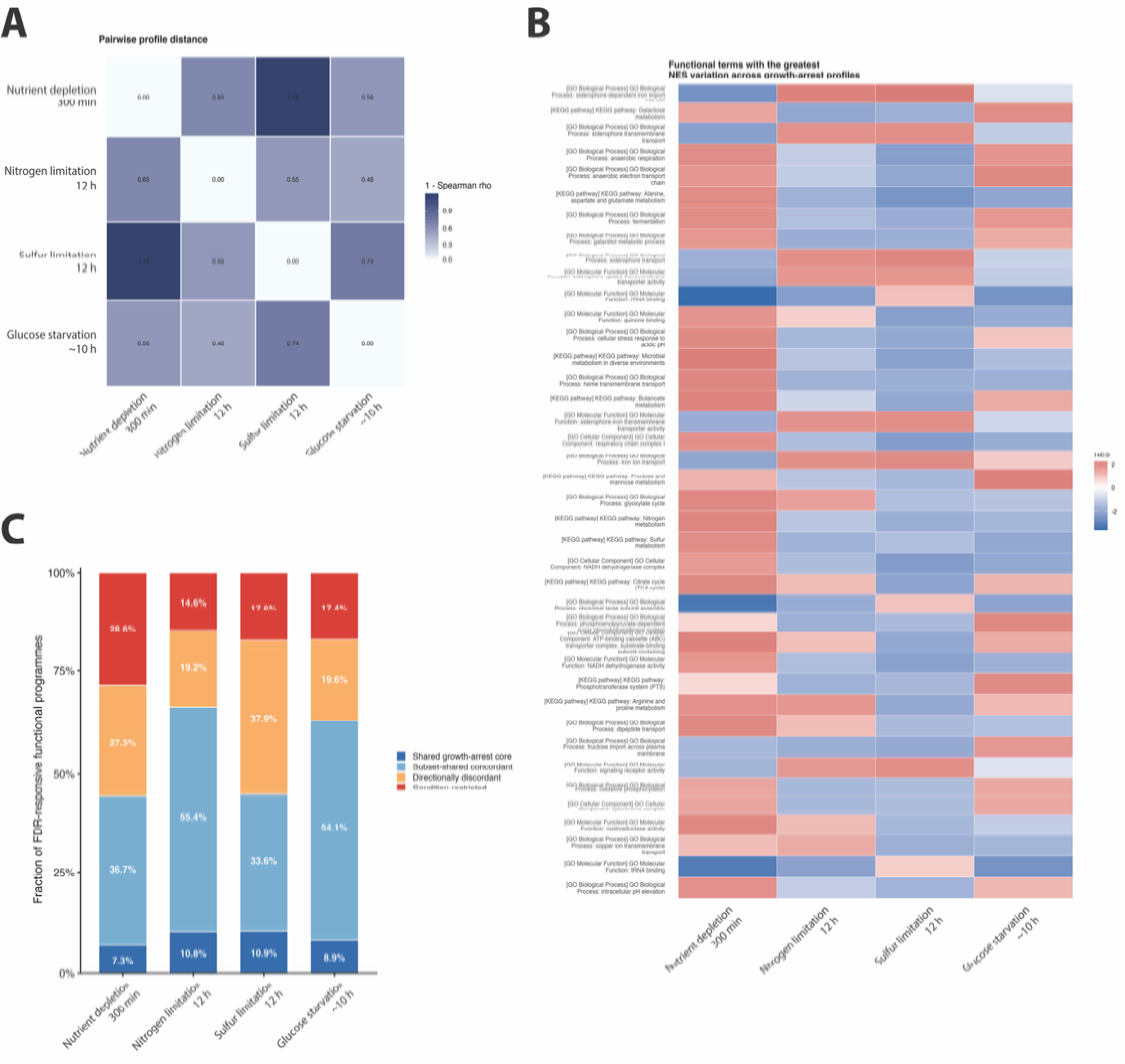

**Supplementary Fig. 9. Distinct transcriptional programmes accompany a shared RNA structural-stability signature across nutrient-limited growth-arrest conditions in *Escherichia coli*.** (**A**) Pairwise Spearman distances (1 − ρ) between complete normalized enrichment score (NES) profiles generated by pre-ranked GSEA across Gene Ontology terms and KEGG pathways. Profiles compare 300-min nutrient depletion with the matched 0-min baseline (GSE224463), 12-h nitrogen or sulfur limitation with the matched 3-h baseline (GSE226643), and established glucose starvation with pre-starvation (GSE131992). (**B**) NES values for the functional programs showing the greatest variation across the four growth-arrest contrasts. Positive and negative values indicate enrichment among transcripts with increased and reduced abundance, respectively, relative to the matched reference state. (**C**) Composition of FDR-responsive programs (BH-adjusted *P* < 0.25) classified as a directionally concordant core shared across all four contrasts (dark blue), subset-shared concordant programs present in two or three contrasts (light blue), directionally discordant programs (light orange), or condition-restricted programs (dark orange-red). Fractions were calculated within each contrast and sum to 100%.
**Supplementary tables**

The supplementary tables supporting this study are accessible through the Zenodo repository:

<https://zenodo.org/records/17363927?preview=1&token=eyJhbGciOiJIUzUxMiJ9.eyJpZCI6IjRiNmY4YTk0LWI5ZWYtNDhjNy05MTRlLTNlYzQ1ZTRiNzE5YiIsImRhdGEiOnt9LCJyYW5kb20iOiJmNjE1YTQxMzkxZWQ3ODVlMDQ1ZDM4ZjhmNGRlZjE0YiJ9.u4SvYzmHKrGQxqc-sREQozT1h1iaX5z7CLcXX1fBRs5dEnsEtLExsxZki201wIDt_WHSqBnsPc8AVlc7KtIkcw>
